# *In silico* analysis of *Bacopa monnieri* (L.) Wettst. compounds for drug development against Neurodegenerative Disorders

**DOI:** 10.1101/2022.03.30.486025

**Authors:** Satyam Sangeet, Arshad Khan, Saurov Mahanta, Nabamita Roy, Sanjib Kumar Das, Yugal Kishore Mohanta, Muthupandian Saravanan, Hui Tag, Pallabi Kalita Hui

## Abstract

Neurotrophins play a crucial role in the development and regulation of neurons. Alterations in the functioning of these Neurotrophins leads to several Neurodegenerative Disorders. Albeit engineered medications which are accessible for the treatment of Neurodegenerative Disorders, due to their numerous side-effects, it becomes imperative to formulate and synthesize novel drug candidates. Plants could be utilized as an alternative for these manufactured medications because of their low incidental effects in contrast with the engineered drugs. *Bacopa monnieri* has been traditionally known to be utilized to treat Neurodegenerative Disorders. Therefore, in current study an *in-silico* based study was carried out to evaluate the pharmacological effect of *Bacopa monnieri*. Molecular Docking was carried out to screen the active phytochemicals of *Bacopa monnieri* which can act as potential drug candidates against the causative proteins of Neurodegenerative Disorders. A total of 105 biologically active phytochemicals from *Bacopa monnieri* were docked against the receptors of brain-derived neurotrophic factor, neurotrophin-3, neurotrophin-4, and nerve growth factor. Based on molecular docking study it was observed that the phytocompounds Vitamin E, Benzene propanoic acid, 3,5-bis(1,1dimethylethyl)4-hydroxy-, methyl ester (BPA), Stigmasterol, and Nonacosane of *Bacopa monnieri* significantly fits to the active residues of the four selected drug targets. Further Molecular Dynamics simulation study was performed to examine the stability of the binding of these phytochemicals with the selected targets. Drug likeness properties as well as related physico-chemical properties were analyzed through ADMETox study. Our findings suggested that the phytocompounds Vitamin E, BPA, Stigmasterol and Nonacosane significantly bind against brain-derived neurotrophic factor, neurotrophin-3, neurotrophin4, and nerve growth factor, respectively which may be the potential drug candidates for the treatment of neurodegenerative disorders.

**Graphic Abstract:** 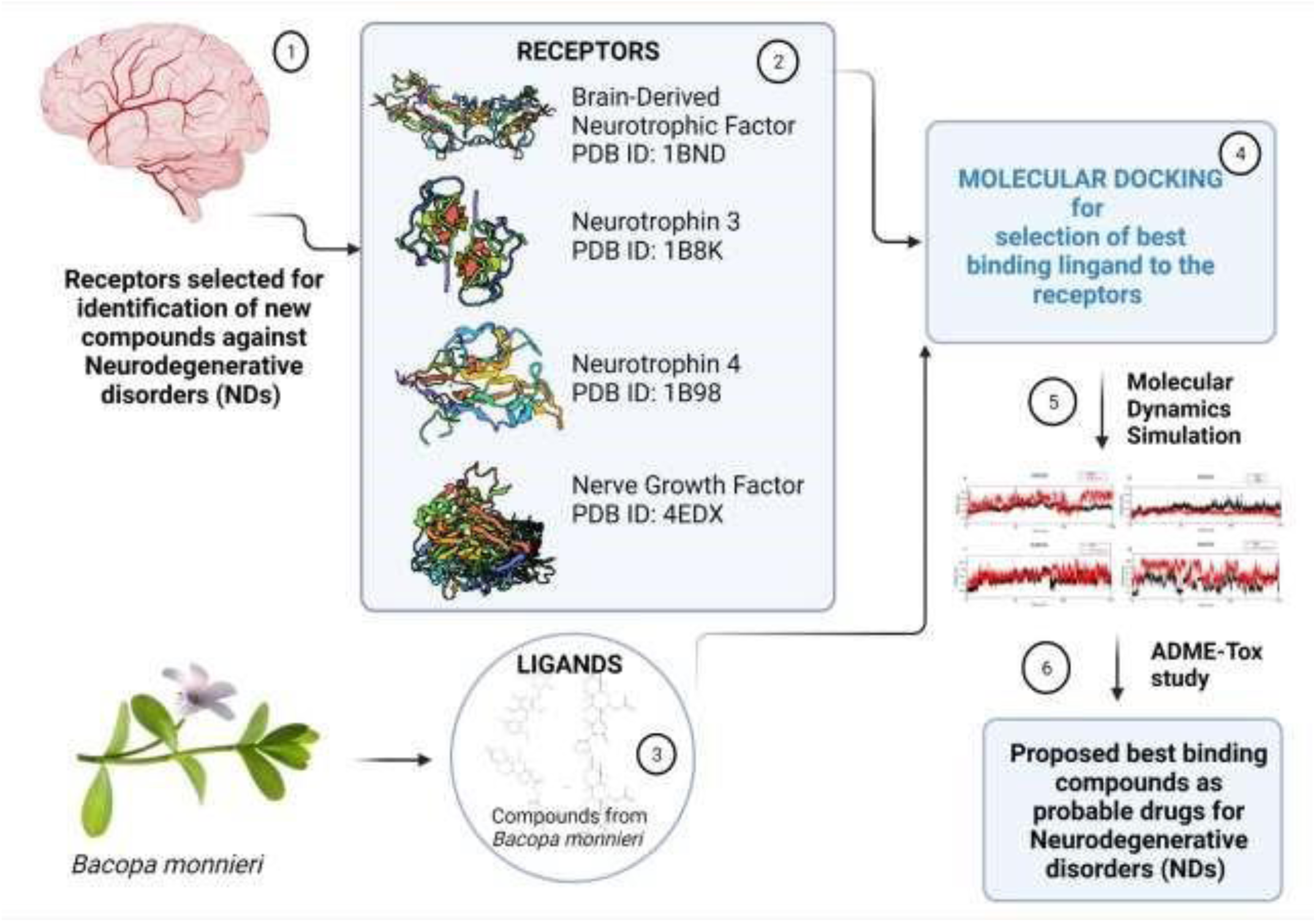

## 1. INTRODUCTION

Neurodegenerative disorders (NDs) are recognized to be noteworthy threats to human health. There is a huge economic burden of NDs on the global, national, and regional levels. Numerous symptoms are associated with NDs including memory loss, tremors, forgetfulness, agitation. Many different mechanisms lead to NDs such as apoptosis, protein aggregation, oxidative stress, cytotoxicity and aging. The NDs have a major involvement of neurotrophins, which are group of proteins having related structures and functions and have an important involvement in survival [1, 2], maintenance [3], function [4, 5], regulation [6], and development [7] of nerve cells. The mammalian cells include four types of neurotrophins: brain-derived neurotrophic factor (BDNF) [8], neurotrophin 4(NT 4/5) [9], nerve growth factor (NGF) [10], and neurotrophin 3 (NT-3) [11]. Neurotrophins belong to a class of growth factors that can regulate cell survival, growth and differentiation. Thus, adequately termed as neurotrophic factors [12]. Neurotrophic factors that mammalian cells secrete play an important role in the survival of nerve cells by preventing them from participating in programmed cell death. Decrease in the level of neurotrophins results in several NDs [13, 14]. Several studies have considered Neurotrophins to be a potential target for the treatment of NDs [15–17].

All the four neurotrophins bind individually by same affinity with two different receptor types, a common neurotrophin receptor P75^NTR^ and a specific receptor of tyrosine kinase Trk family, and thus activates the signal cascade response [18]. P75^NTR^ receptor is structurally related to the tumor necrosis factor receptors (p55^TNFR^ and p75^TNFR^). Involvement of the Arg88 residue of BDNF has been shown to actively participate while interacting with the P75^NTR^ receptor. In NGF, residue Lys32 has been found to be actively associated with the P75^NTR^ binding. Also, the involvement of residues Trp21, Ile31, Lys88, and Arg100 with the binding of P75^NTR^ has been reported. Interaction of NT3 with P75NTR is guided by the involvement of Gln83 and Arg103. NT4 shows the major involvement of Arg98 which actively binds to P75^NTR^ [18].

While the binding of all the above three neurotrophins with p75NTR results in the transmission of neurotrophic signals, binding of NGF to p75^NTR^ can initiate a signaling process leading to either survival (NF-***κ***B pathway) or apoptosis (Rac1/JNK pathway) of nerve cells. Kuner and Hertel [19] showed that the binding of NGF to p75^NTR^ induces an apoptotic cycle in the neuroblastoma cell lines making it imperative to understand the binding behaviour of NGF with p75NTR and find potential small molecules to inhibit their interactions. Colquhoun et al. [20] demonstrated that a compound PD90780 [7-(benzolylamino)-4,9-dihydro-4methyl-9-oxo-pyrazolo[5,1-b]quinazoline2-carboxlic acid] interacts with NGF and inhibit its binding to p75^NTR^ thereby halting the process of apoptosis. These fragmented pieces of evidence make it imperative to understand the potential interaction between neurotrophins and p75ntr.

An Indian medicinal plant *Bacopa monnieri* (BM) has been reported to impart healing against neurodegenerative disorders since long time. Several standard extracts of BM have shown improvement in many neurodegenerative disorders [21–24]. BM is well recognized to have a potential in rejuvenating nerve cells and improving cognition as well as memory [25]. A recent study proved that *B. monnieri* extract enhanced the cognition effectively [22]. Further studies also identified the various bioactive components contained by BM such as saponins, triterpenoid bacosaponins and saponins glycosides which actually provide the healing in NDs and improve memory power. For example, a saponin Bacoside A is reported to improve memory power [26]. Also, when extracting aerial parts of BM administered orally, it produces a neuroprotective effect in cold stress induced hippocampal neurodegeneration of rats [27].

Thus, the current study focuses on understanding the interaction between neurotrophins and p75ntr and explores the potential of phytochemicals of BM as potent inhibitors against the interactions of neurotrophins and receptors. In-silico evaluation was undertaken to screen out the phytochemicals of BM which inhibits the binding of neurotrophins (BDNF, NT3, NT4, and NGF) to p75^NTR^ through molecular docking, molecular dynamics simulations followed by ADMET analysis to explore the clinical relevance of the compounds.

## 2. Materials and method

### 2.1. Selection of phytochemicals for the study

*B. monnieri* contains many phytocompounds that are belonging to the class of saponins, triterpenoid bacosaponins and saponins glycosides [26]. Among these phytocompounds more than 100 biologically active phytochemicals were selected for the study which were collected from published literature and also Dr Duke’s Phytochemical and Ethnobotanical Database [28]. The phytochemicals were listed out in order to download their structure.

### 2.2. Ligand and Receptor Preparation

The 3-D structures of selected phytocompounds (*sdf format*) and their canonical SMILES were downloaded from PubChem Compound (https://pubchem.ncbi.nlm.nih.gov/) database [29] and converted to PDB format using Open Babel software (http://openbabel.org/) [30]. The 2D structure of those phytochemicals whose 3D structures were not available in PubChem database were downloaded and converted into 3D using Open Babel software. The energy minimizations was carried out using Chem3D software (ChemOffice, 2002) [31] through MM2 minimization (Section S1).

For preparation of the receptor structures, the PDB structures of Brain-Derived Neurotrophic Factor (PDB ID: 1BND), Neurotrophin 3 (PDB ID: 1B8K), Neurotrophin 4 (PDB ID: 1B98) and Nerve Growth Factor (PDB ID: 4EDX) receptors were downloaded from RCSB Protein Data Bank (www.rcsb.org). The receptors were prepared by deleting all the water molecules and unwanted residues followed by energy minimization through Chimera software [32]. PyMOL (Schrodinger, 2010) was used for the visualization of the 3D structures of both ligands and receptors in the present study [33]. The p75NTR acts as a common natural ligand for all the four receptors BDNF, NT-3, NT-4 and NGF. The structure of p75NTR (PDB ID: 3BUK) was also downloaded from RCSB Protein Data Bank and energy minimized using Chimera.

### 2.3. Molecular Docking study

PatchDock online server (https://bioinfo3d.cs.tau.ac.il/PatchDock/) was employed for the molecular docking studies between the receptors (PDB ID: 1BND; 1B8K; 1B98; 4EDX) and the ligands (phytochemicals) [34, 35]. The active sites of the receptors served as the ligand binding site, which were revealed through molecular docking of receptors (BDNF, BT3, NT4 and NGF) with their common natural ligand p75NTR. The clustering RMSD values were set at default (4.0 Å) as suggested by the server before the start of molecular docking. The respective Neurotrophin structures were uploaded in the Receptor molecule option and the phytochemical structures were uploaded in the Ligand Molecule option and molecular docking was performed. The PatchDock score and the ACE value were tabulated and the result was visualized in Maestro (Schrodinger 2020).

### 2.4. Molecular Dynamics Simulations

MD simulations were performed to analyze the binding behavior of the phytochemicals to the selected Neurotrophins in order to study the dynamic behavior of the protein-ligand interactions. The MD simulation was carried out using GROMACS version 2018.1 for 150ns (nanosecond) [36– 38]. CHARMM36 all-atom force field and TIP3P water model were used to generate the protein topology parameters. CGenFF (CHARMM General Force Field) server was used to generate the ligand topology. The ligand-protein complex obtained from docking studies were then solvated in a cubic box with TIP3P water model. Temperature of the system was gradually stabilized from 0K to 300K for 150ns under NVT ensemble. The system was then simulated under NPT ensemble at 300K temperature and 1.0 bar pressure. The 150ns production run was carried out and the system coordinates were saved after every 20ps for post processing analysis.

#### 2.4.1. Root Mean Square Deviation (RMSD)

The structure and dynamic properties of the protein-ligand complexes were analyzed as the backbone RMSDs during the simulation period of 150 ns [39]. The RMSD was measured as the average distance between the backbone atoms of the protein-ligand structures and it was derived from the following equation:

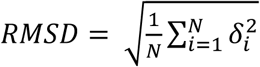

Where N represents the total number of atoms considered in the calculation and δ represents the distance between the N pairs of equivalent atoms.

#### 2.4.2. The root mean square fluctuations (RMSF)

The root mean square fluctuations (RMSF) were assessed and plotted to equate the flexibility of each residue in the–ligand-protein complexes. The RMSF of the protein ligand complex denoted the minimized fluctuation for all the complexes [39]. Python package, Matplotlib, was used for plotting the RMSF curves [40].

#### 2.4.3. Radius of Gyration (RoG)

The radius of gyration (RoG) was calculated for the respective Neurotrophin backbone. Radius of gyration of a protein is the measure of its compactness. If a protein is stably folded, its corresponding radius of gyration will have a relatively stable value of Rg. On the other hand, if the protein is unstable or it starts to unfold, its radius of gyration will be reflected by a fluctuating value of Rg.

#### 2.4.4. Total Energy

The total energy of the system after the MD Simulation shows the total stability and distribution of the system throughout the simulation time. If the protein-ligand system is steady, the total energy curve of the system will remain stable throughout the simulation.

#### 2.4.5. Drug Likeness screening of ligands

The phytochemicals were also screened against the commercial drugs by calculating their Lipinski’s value (http://www.scfbio-iitd.res.in/software/drugdesign/lipinski.jsp) [41].

Lipinski’s Rule of Five (RO5) is set of factors that allow the early screening of phytochemicals compared to the commercial drugs.

### 2.5. ADME-tox Studies

Adsorption, Digestion, Metabolism, Excretion and Toxicity studies were performed using SWISS-ADME (http://www.swissadme.ch/) [42] and Pre-ADMET server (https://preadmet.bmdrc.kr) [43]. The structure of drugs and phytochemicals were uploaded and the corresponding drug likeness features such as lipophilicity, solubility and drug likeness score were obtained.

## 3. Results

### 3.1. Selection of phytochemicals for the In-silico study

100 biologically active phytocompounds were selected for the study that were referred from previous literature [28]. The top hit phytochemicals are presented in Table 1. The rest of the phytochemicals are listed in Table S7. Their structures, molecular weights were verified from PubChem (https://pubchem.ncbi.nlm.nih.gov/) database [29]. We identified top-hit phytochemicals for each neurotrophins and performed molecular dynamic simulations to analyze the binding behavior of the compounds with neurotrophins. The result shows a better binding behavior of the phytochemicals with the neurotrophins suggesting that these small molecules from the medicinal plants can prove a potential drug candidate against neurodegenerative disorders.

**Table 1:** Molecular docking of phytocompounds from *B. monnieri* against the selected receptors.

| Neurotrophins | Phytochemicals | PatchDock Score | Atomic Contact Energy (kcal/mol) | Interacting Residues |
| --- | --- | --- | --- | --- |
| BDNF | Vitamin E<br>PubChem CID: 14985 | 4592 | -246.62 | Trp19, Tyr54, Phe53, Tyr52, Gln51, Lys50, Leu49, Val42, Val44, Val87, **Arg88**, Ala89, Leu90, Trp100, Ile105 |
| NT3 | Benzene propanoic acid, 3,5-bis (1,1-dimethylethyl)-4hydroxy-, methyl ester<br>PubChem CID: 62603 | 3630 | -26.71 | Leu19, Val21, Lys24, Ser26, Ala27, Ile28, Gln34, Arg56, Ile104, **Gln83, Arg103**, Asp105, Glu54 |
| NT4 | Stigmasterol<br>PubChem CID :5280794 | 5684 | -409.31 | Gly48, Ala47, Ala46, Val44, Trp110, Trp112, Gln54, Phe56, Val97, **Arg98**, Ala99, Tyr96, Leu52, Tyr55, |
| NGF | Nonacosane<br>PubChem CID: 12409 | 5736 | -171.43 | Gln95, Ala97, Trp99, Arg100, Phe101, Lys88, Val20, Ala98, **Lys32, Ile31**, Phe54, **Trp21**, Ser19, Val18, Arg59, Ser17, Phe49 |

*\*The interacting residues of the active site are highlighted in bold*

### 3.2. Ligand and Receptor preparation

The structure of the receptors is represented in Fig1. The active site of the receptors is highlighted in yellow. Arg88 has been shown to actively interact with p75NTR receptor. NT3 interacts with p75NTR via two important residues, Gln83 and Arg103. NT4 shows a major involvement of Arg98 while interacting with p75NTR receptor. Interaction of NGF is well guided via residue Lys32, but it has also been seen that the involvement of residue Trp21, Ile31, Lys88 and Arg100 also guide a better interaction with p75NTR receptor [18]. The structure of the respective Neurotrophins with their active site residues are represented in Fig1. The crystal structure of the Neurotrophins is shown in Fig.S1.

**Figure 1:**
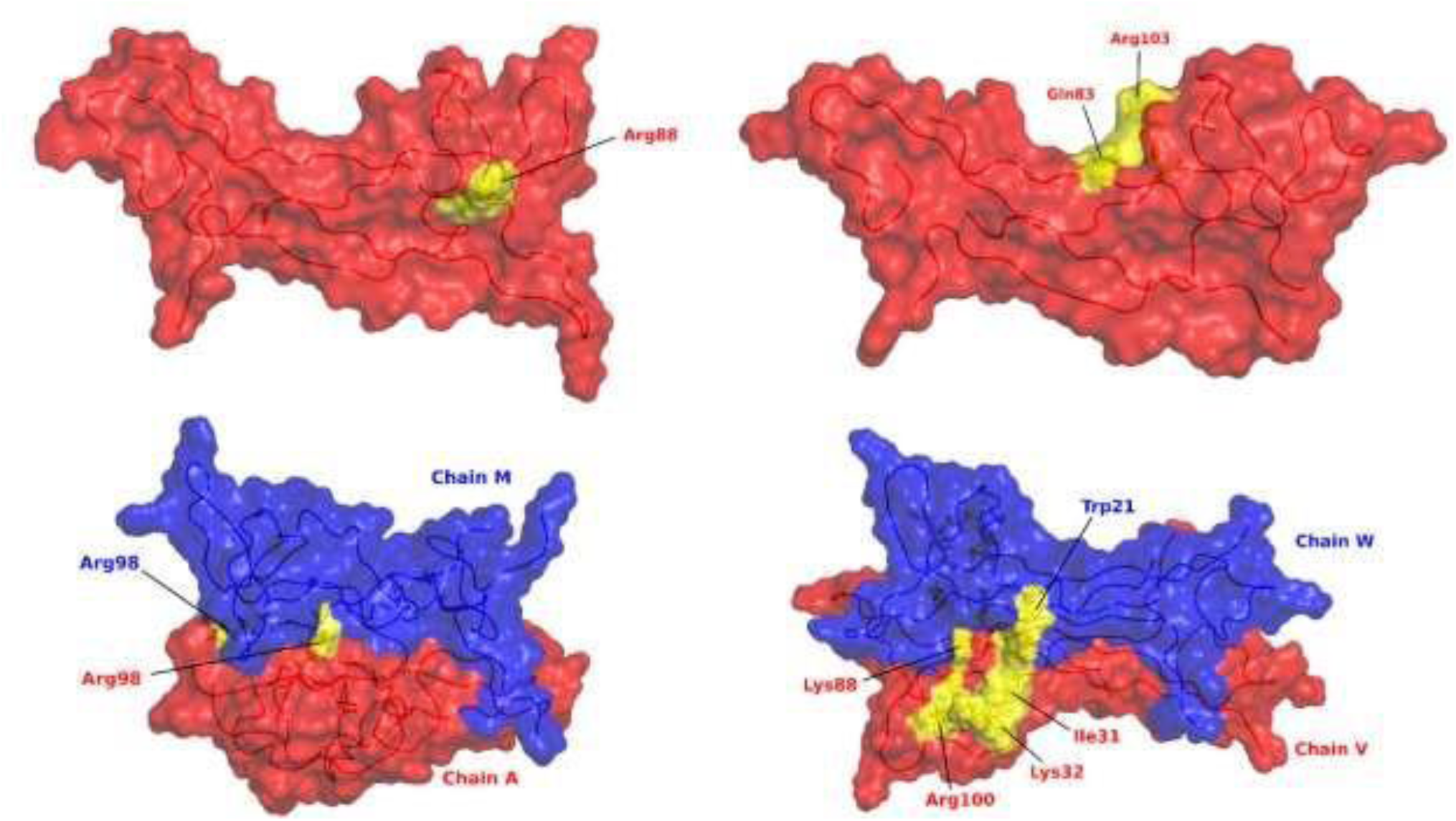
Surface view of the Neurotrophins showing the active site residue highlighted in yellow. (a) BDNF with the active site residue Arg88 in yellow highlight (b) NT3 with the active site residue (Gln83 and Arg103) (c) NT4 Homodimer. Chain A is in red color and Chain M is in blue color. The active site residues are highlighted in yellow while their labelling is done with respective color to show the residue in chain wise manner (d) NGF Homodimer. Chain V is in red color and Chain W is in blue color. The active site residues are shown in yellow highlight. A similar active site is also present on chain W which is not visible due to the 2D image. The residues are colored in order to show the native chain to which they belong.

### 3.2. Molecular Docking

PatchDock online server (https://bioinfo3d.cs.tau.ac.il/PatchDock/) was employed for the molecular docking studies between the receptors (PDB ID: 1BND; 1B8K; 1B98; 4EDX) and the ligands (phytochemicals). The binding affinity between the receptor and ligand was determined based on PatchDock score. Higher PatchDock score indicates better binding affinity. Out of the 26 phytochemicals docked against the four selected receptors PDB ID: 1BND; 1B8K; 1B98; 4EDX. Vitamin E showed highest interaction against BDNF with a docking score of 4,592 and an Atomic Contact Energy of -246.62 kcal/mol and also showed hydrophobic interactions with Val87, Ala89,

Leu90, Trp100 followed by positively charged interactions with Arg88 and Lys50 (Figure 2A). Similarly, phytocompounds Benzene propanoic acid, 3,5-bis(1,1-dimethylethyl)-4hydroxy-, methyl ester showed highest binding affinity against NT3 with a docking score of 3,630 and an Atomic Contact Energy value of -26.71 kcal/mol followed by hydrophobic interactions with Leu19, Val21, Ala27, Ile28 and Ile104. It also showed a polar interaction with Gln83, Gln34 and Ser26 along with a positively charged interaction with Arg56, Arg103 and Lys24. Interaction with Glu54 and Asp105 was also observed via a charged negative interaction (Figure 2B). However, more than 10 numbers of phytocompounds showed significant interaction against NT4 and NGF. Among these phytocompounds, Stigmasterol showed highest binding affinity towards NT4 with a docking score of 5,684 and an Atomic Contact Energy of -409.31 kcal/mol (Figure 2C). It showed hydrophobic interactions with Ala47, Ala45, Val44, Trp110, Trp112, Phe56, Val97 and Ala98 along with a polar interaction with Gln54 and a positively charged interaction with Arg98. For NGF, Nonacosane showed significant binding affinity with a docking score of 5,736 and an Atomic Contact Energy value of -171.43 kcal/mol (Figure 2D) followed by hydrophobic interactions with Trp21 and Ile31 along with a positively charged interaction with Lys32 of active sites. The interacting residues for respective Neurotrophins are represented in Table-1.

**Figure 2:**
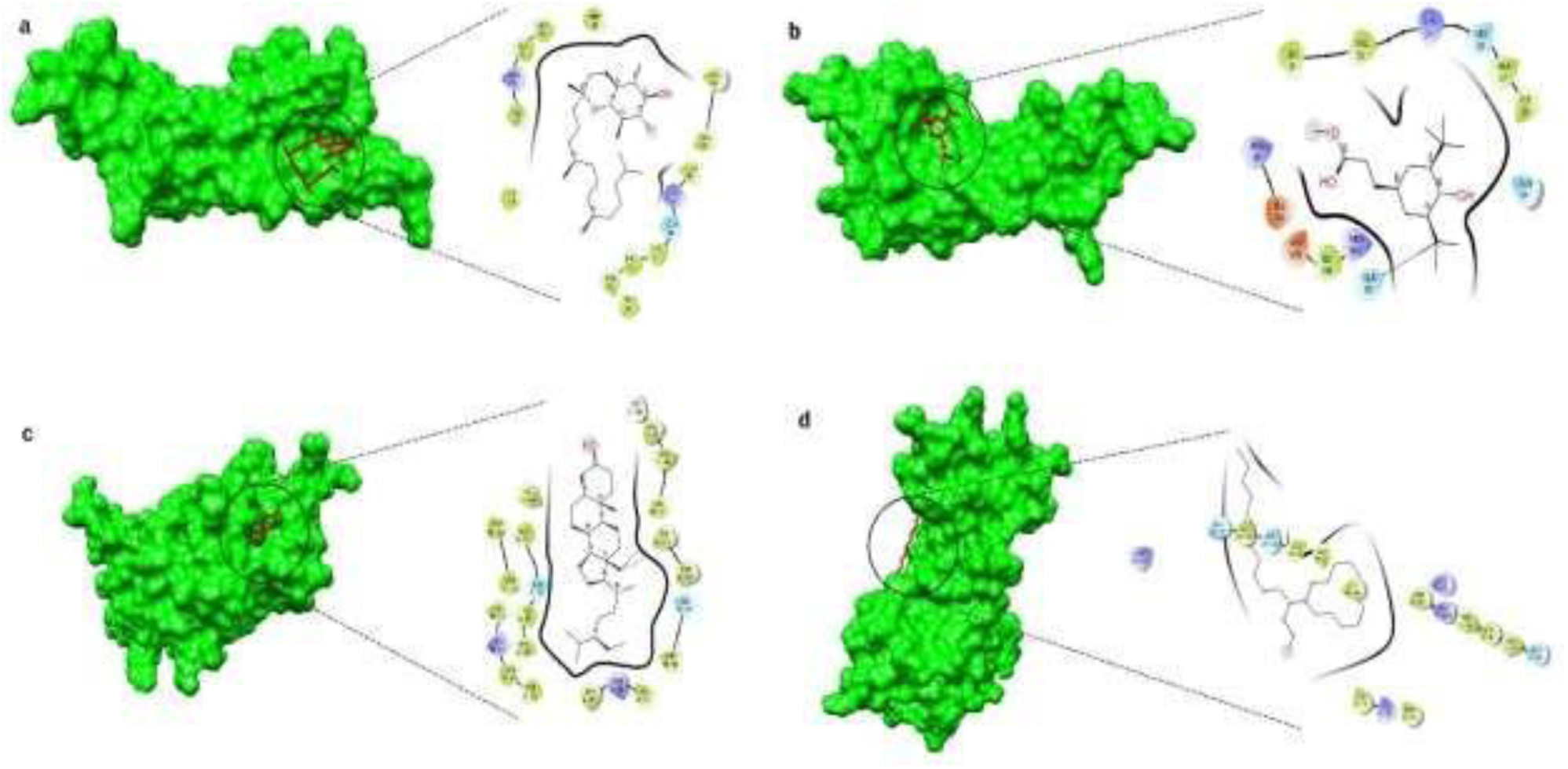
Interaction of a) BDNF (green) and Vitamin E (red); b) NT3(green) and Benzene propanoic acid, 3,5bis(1,1dimethylethyl)-4hydroxy-, methyl ester (red); c) NT 4 (green) and Stigmasterol (red) ; d) NGF (red) and Nonacosane (green). The corresponding interactions of residues with their respective Neurotrophins are represented by the corresponding color codes: Red (Charged negative), Blue (Charged positive), Green (Hydrophobic), Sky blue (Polar), White (Glycine)

The details of interaction of other prominent phytochemicals with their respective Neurotrophins are given in Supplementary Information (Table S6).

### 3.3 Molecular Dynamic Simulations

Molecular simulations of the protein-ligand complexes were subjected to a 150ns MD simulation. The corresponding RMSD, RMSF, Radius of Gyration and Total Energy parameters were calculated to analyze the docking results of the top phytochemical interaction of individual Neurotrophins, i.e., Vitamin E with BDNF, Benzene propanoic acid, 3,5-bis-(1,1dimethylethyl)-4-hydroxy-, methyl ester with NT3, Stigmasterol with NT4 and Nonacosane with NGF. Throughout the simulation time of 150ns, the ligand conformations were extracted corresponding to 30 ns, 50 ns, 100 ns and 140 ns (Fig. 3). These configuration shows the evolution of the ligand throughout the simulation process exhibiting that the ligand explores the potential configurational space to adapt to the most stable configuration complimentary to the binding pocket of the respective neurotrophins. The Free Energy Surface (FES) profile of the most stable configurations for the respective ligand is shown in the Supplementary figure S2.

**Figure 3:**
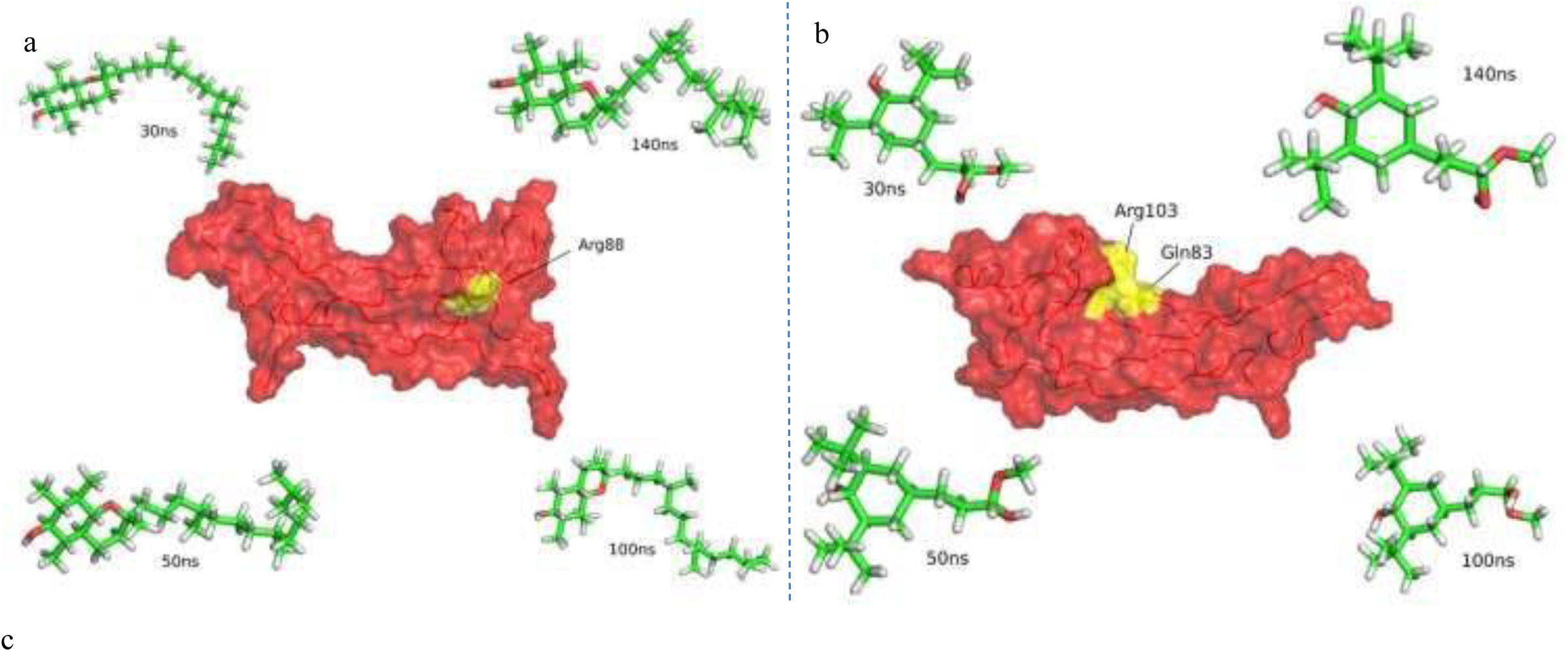
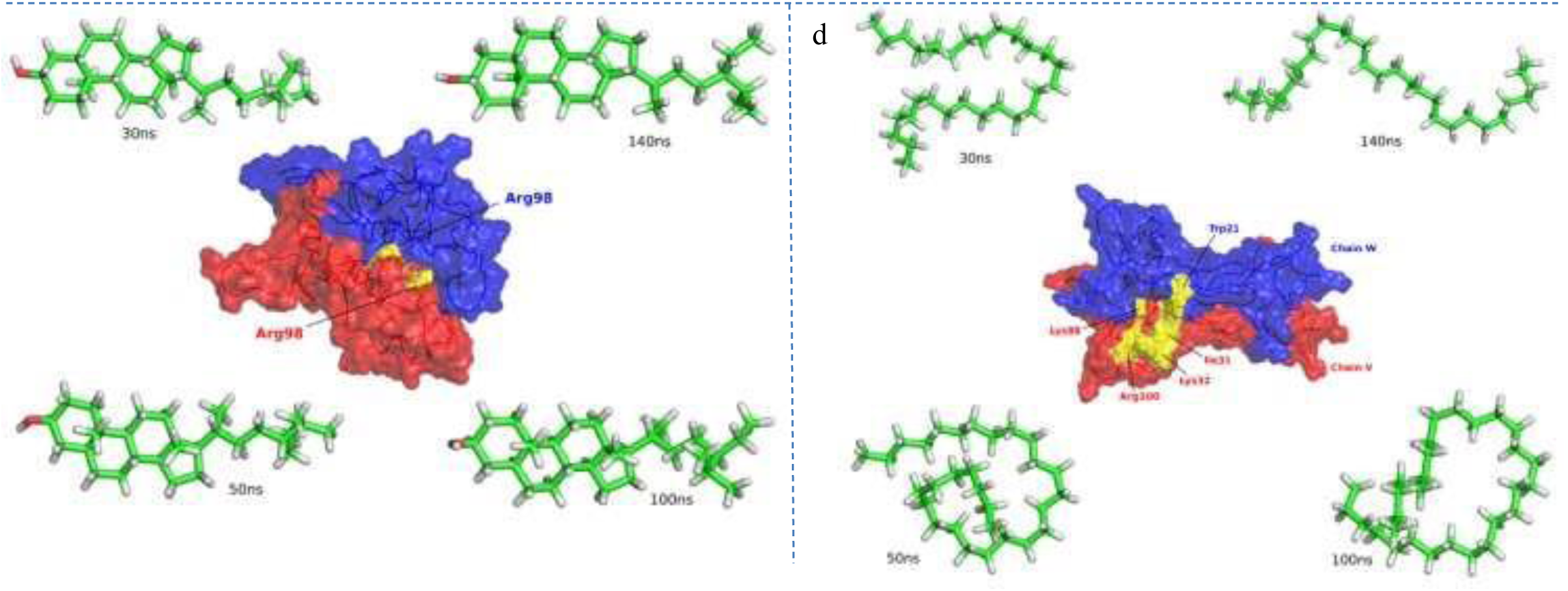
Structure of Neurotrophins showing the active site (highlighted in yellow) and four configurations of the ligand is shown corresponding to different time frames during the Molecular Dynamic Simulation run of 150ns. (a) BDNF with active site highlighted in yellow and ligand (Vitamin E) configurations (b) NT3 with active site highlighted in yellow and ligand (BPA) configurations (c) NT4 Homodimer [Chain A: red & Chain M: blue] with active site highlighted in yellow and ligand (Stigmasterol) configurations (d) NGF Homodimer [Chain V: red & Chain W: blue] with active site highlighted in yellow and ligand (Nonacosane) configurations.

#### 3.3.1. Root Mean Square Deviation (RMSD)

Root Mean Square Deviation (RMSD) denotes the stability of the protein-ligand complex throughout a MD simulation. BDNF, NT3 and NT4 Neurotrophins remained stable throughout the 150ns MD Simulation (Fig.4). The RMSD fluctuations were in the admissible limit of 0.1-0.3 nm, 0.1-0.4 nm, 0.05-0.25 nm for BDNF (Fig 4A), NT3 (Fig 4B), NT4 (Fig 4C). NGF exhibited a slightly higher fluctuation within 0.1-0.6 nm (Fig 4D) compared to other Neurotrophins. The RMSD of Vitamin E (Fig.4a) shows that the binding between the ligand and receptor got stabilized after 120ns resulting in a stable fluctuation in the range 0.2-0.3nm and 0.35-0.5nm for BDNF and Vitamin E respectively. The binding between N3 and BPA remained stable throughout the simulation time (Fig.4b) with average fluctuation ranging from 0.2-0.4nm and 0.1-0.2nm for N3 and BPA respectively. NT4 and Stigmasterol showed a stable binding till 80ns after which the fluctuation starts to increase. The fluctuation in the end tail of Stigmasterol might be a possible cause for the rapid fluctuation post 80ns. On the other hand, NGF remained fluctuating throughout the simulation time (Fig.4d). The linear structure of Nonacosane might be a possible cause for the unregulated fluctuation emerging in NGF. Although the ligand acquires different structural conformations throughout the simulation in order to assist in a better binding affinity, but the fluctuation in the NGF’s backbone remained high throughout. The structure of the ligand with their respective receptor at the most stable free energy change is shown in Fig.S2.

**Figure 4:**
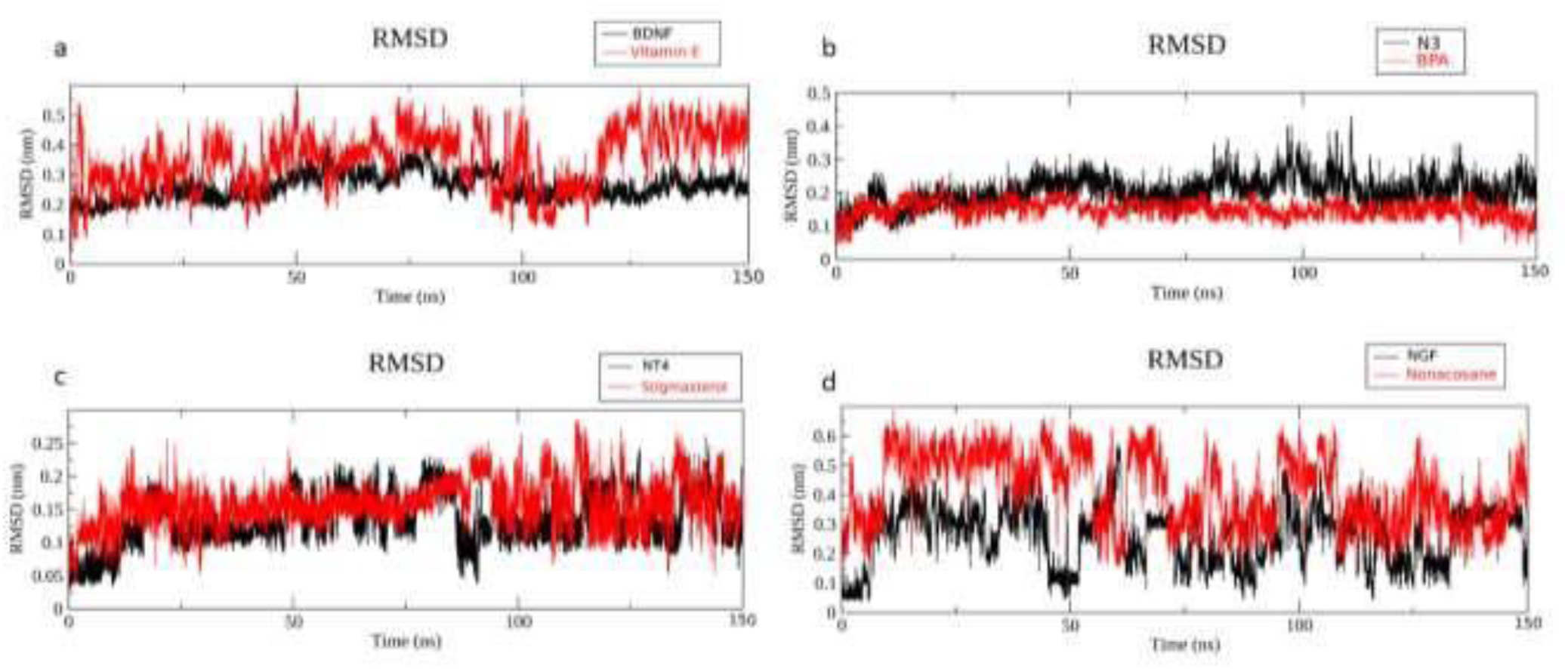
Root Mean Square Deviation (RMSD) of receptors with respective top hit phytochemicals for a Molecular Simulation run of 150ns. (a) RMSD plot of BDNF with Vitamin E (b) RMSD plot of NT3 with BPA (c) RMSD plot of NT4 with Stigmasterol (d) RMSD plot of NGF with Nonacosane Figure 3 represents the different configuration of the ligands with respective receptors throughout the 150ns molecular dynamics simulation. Four structures were taken in in order to show the structural evolution of the ligand. In the case of BDNF (Fig3a), Vitamin E was initially less torsional resulting in a slightly high fluctuation in the starting (Fig.4a). As the time progressed further, Vitamin E took on a linear configuration (Fig.3a 50ns) ultimately achieving a bent structure (Fig.3a 140ns) resulting in a stabilized interaction with BDNF. The binding interaction of BPA with N3 was almost stable throughout the simulation (Fig.4b). Thus, a very little structural evolution was observed for BPA at different time stamps (Fig.3b). Stigmasterol showed a stable binding before 100ns (Fig.4c) which is relevant from the structural evolution of Stigmasterol. The structure of Stigmasterol remained almost the same at 30ns and 50ns (Fig.3c), but after it crosses the 100ns time stamp, the tail end of Stigmasterol starts to wind, almost resembling the pattern of the motion of a propeller (Fig.3c 100ns, 140ns). The slightly higher fluctuation of Stigmasterol post 100ns time stamp might be due to the rotating nature of tail end. As compared to binding characteristic of other phytochemicals, Nonacosane showed a less stable binding with NGF. The RMSD fluctuation of Nonacosane (Fig.4d) was slightly higher indicating that the ligand fluctuates relatively high in order to achieve the right binding conformation. This is evident from the Nonacosane’s structural evolution (Fig.3d) as it achieves different configuration at each time stamp. The initial higher fluctuation might be caused due to the linear nature of Nonacosane ultimately making the ligand change its structural form to achieve a bent down structure around 100ns.

#### 3.3.2. Root Mean Square Fluctuation (RMSF)

The Root Mean Square Fluctuation (RMSF) exhibits the flexibility at residual level. Low level of fluctuations was observed in all the Neurotrophins. The RMSF fluctuation plots reveal the stable behavior of the functionally important residues in respective Neurotrophins (Fig 5). The RMSF studies of Vitamin E (Fig. 5A) suggest that the curve almost fits closely to that of BDNF along with its natural ligand. Looking at the molecular simulation curve of NT3 with the respective phytochemicals it can be seen that Benzene propanoic acid, 3,5-bis(1,1-dimethylethyl)-4hydroxy-, methyl ester (Fig 5B BPA) shows better results as compared with 3A(1H)-Azulenol,2,3,4,5,8,8A-hexahydro-6,8Adimethyl-3-(1-M (Fig. 5B 3A(1H)).When the active amino acid is looked upon on the X-axis, it can be seen that the BPA curve for Gln83 has a dip which goes beyond the curve of NT3 suggesting that BPA has more stable binding with NT3. A similar result can be seen for Arg103 as well. RMSF curve of NT4 with the respective phytochemicals suggest that Stigmasterol (Fig. 5C - yellow) shows better stability while binding with NT4. Looking closely at the active amino acid residue Arg98, the RMSF curves for Stigmasterol is lower than NT4. The molecular simulation studies of NGF with the respective phytochemical shows that Nonacosane (Fig. 5D - blue) showed better results as compared with other phytochemicals. Looking at the RMSF curve of Nonacosane, it can be observed that the curve at amino acid residues Trp21, Ile31 and Lys32 is below the curve of NGF (Fig. 5D, black).

**Figure 5:**
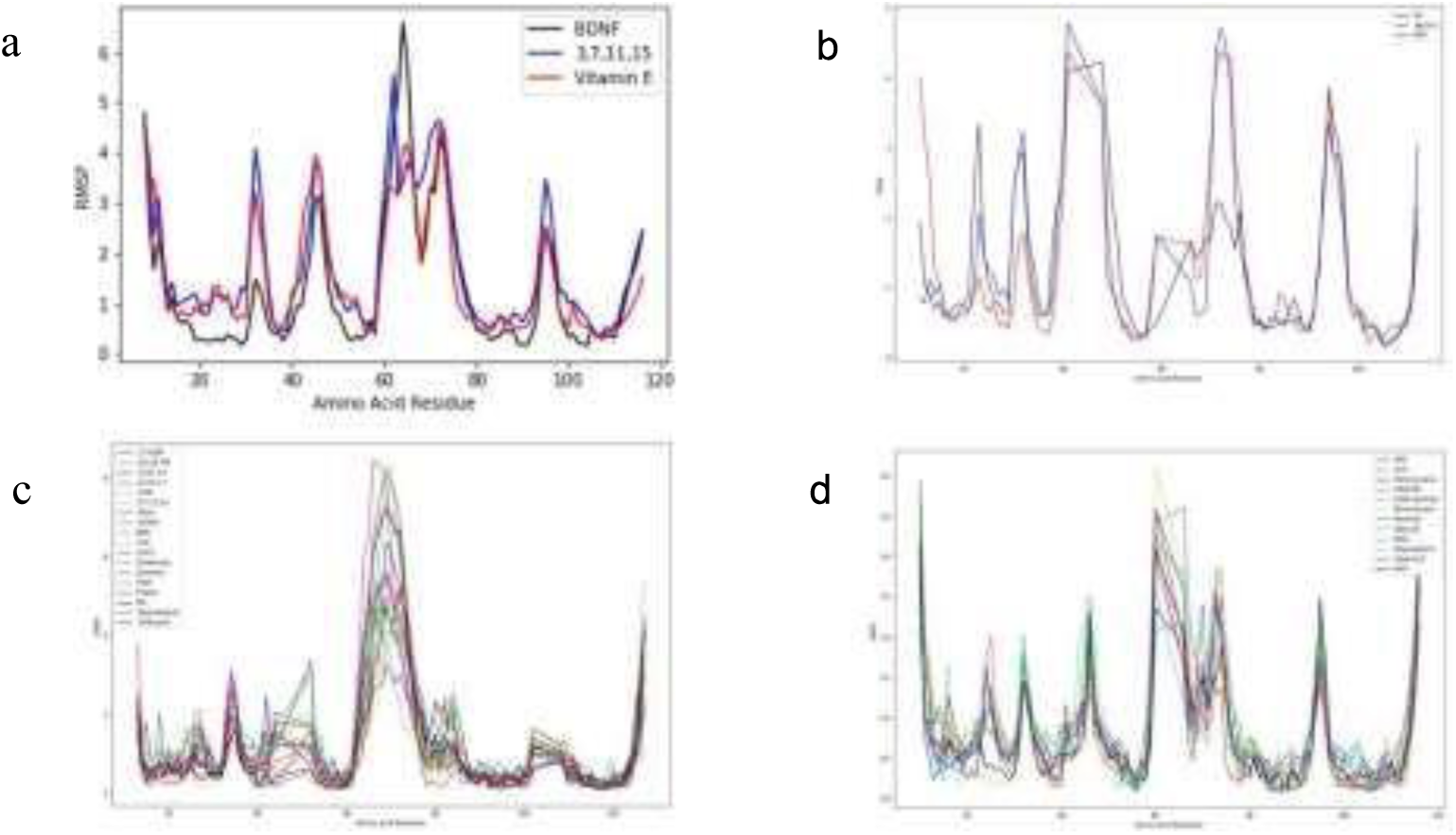
Root Mean Square Fluctuation (RMSF) of (A) BDNF (B) NT3 (C) NT4 (D) NGF

#### 3.3.3. Radius of Gyration (RoG)

The Radius of Gyration represents the protein folding, stability and compactness. The results suggest that BDNF, NT3 and NT4 remained compact and stable throughout the 150ns simulation (Fig 6A, 6B, 6C). The radius of gyration for NGF (Fig 6D) showed a non-uniform behavior between 5-10 Å suggesting that NGF’s compactness was affected by the Nonacosane’s linear structure.

**Figure 6:**
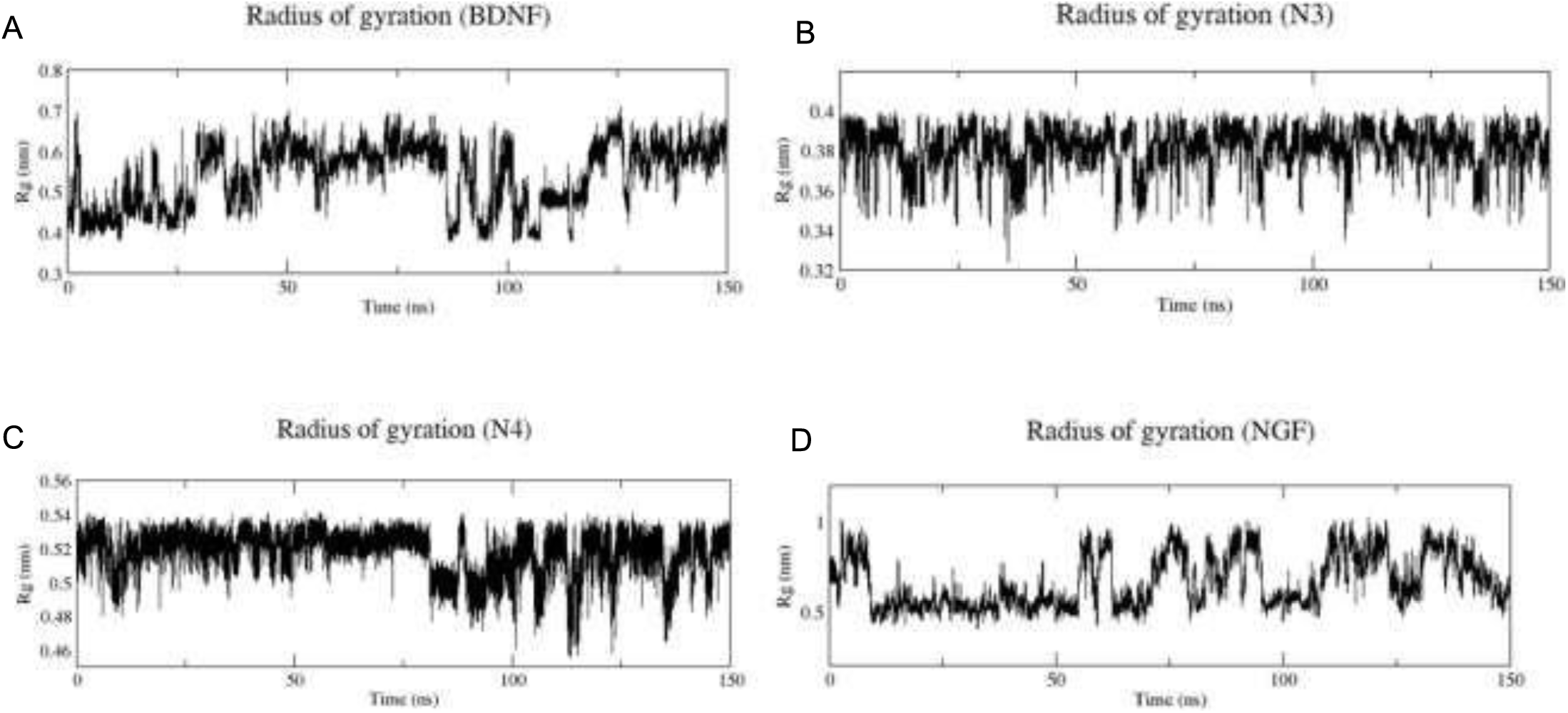
Radius of Gyration (RoG) of (A) BDNF (B) NT3 (C) NT4 (D) NGF

#### 3.3.4. Total Energy

The total energy represents the system’s overall energy stabilization and suggests that the system is energetically minimized. The Total energy of the protein-ligand docked system remained stable throughout the simulation for all the Neurotrophins as represented in Fig 7.

**Figure 7:**
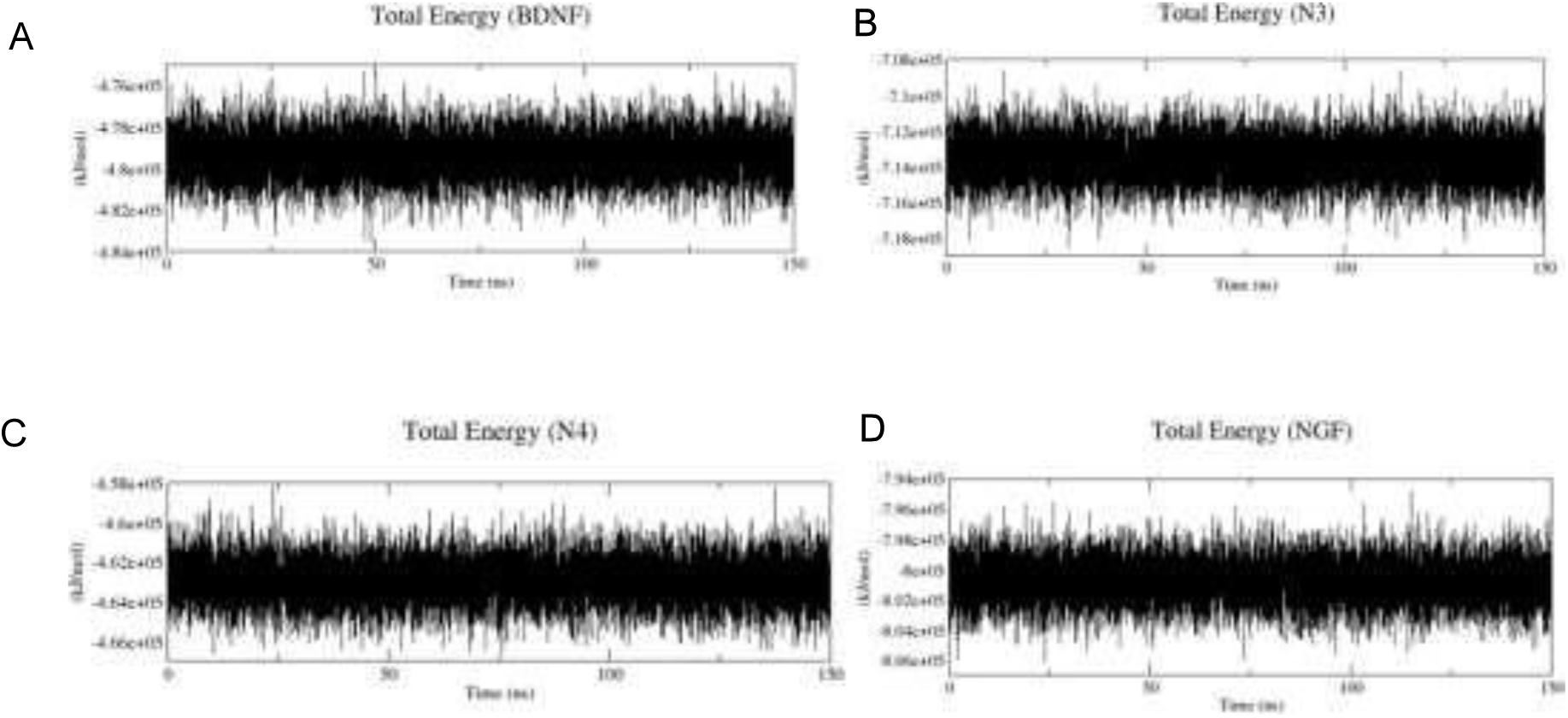
Total energy of (A) BDNF (B) NT3 (C) NT4 (D) NGF

### 3.4. ADME –tox Studies

Drug likeness is the property of any compound to have the potential to act as a drug. Using the online

Lipinski’s Rule of Five (RO5) (scfbio-iitd.res.in/software/drugdesign/Lipinski.jsp) (Table 2), SWISSADME server (https://www.swissadme.ch/) [42] and PreADMET server (https://preadmet.bmdrc.kr/), we analyzed the drug likeness ability and the toxicity of the top hit phytochemicals from *B. monnier*i (Table 3). The detailed methodology of individual studies is given in supplementary information.

**Table 2:**
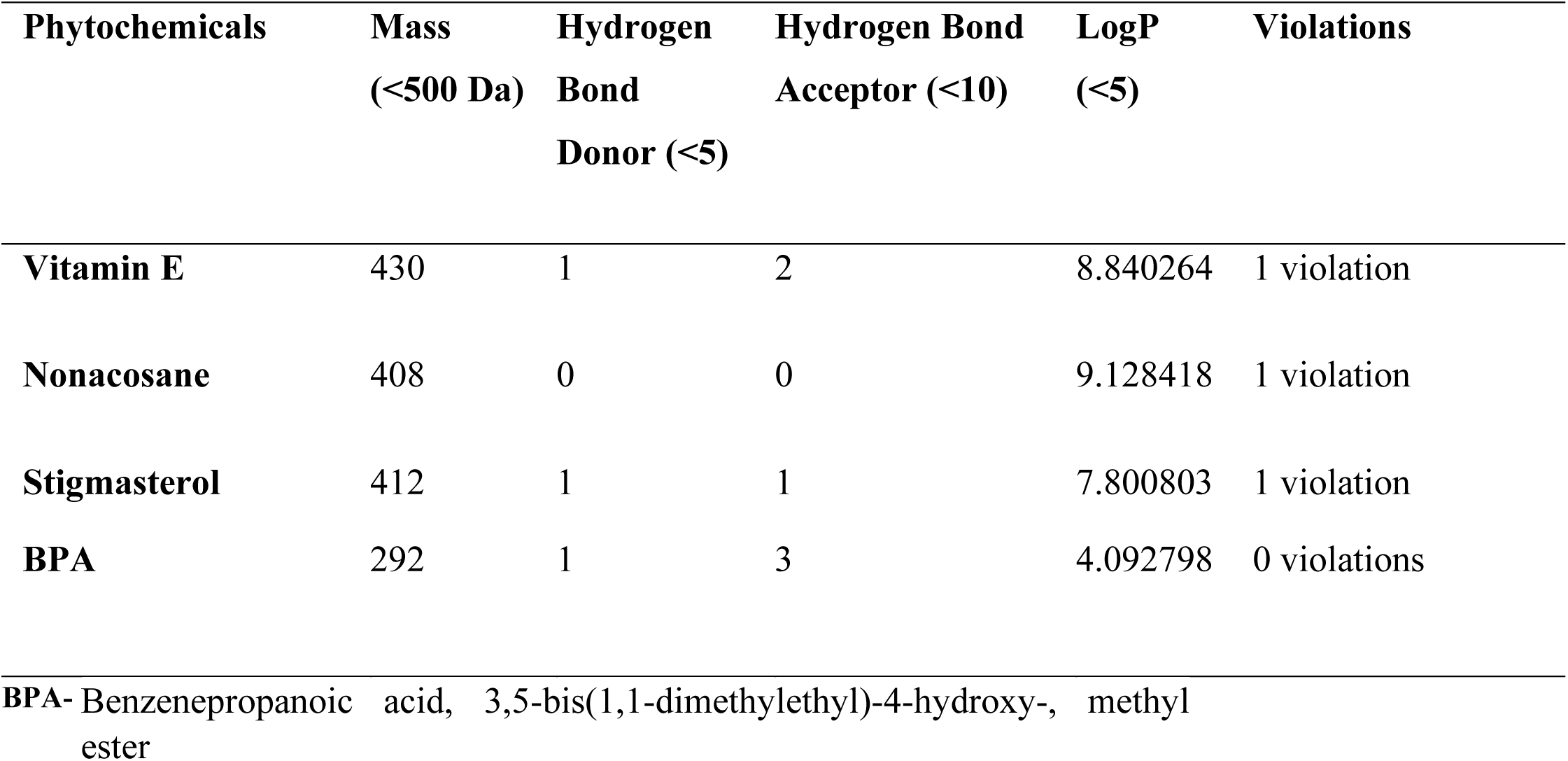
Lipinski’s Rule of Five (RO5) for top hit phytochemicals and Commercial drugs.

| Phytochemicals | Mass<br>(<500 Da) | Hydrogen<br>Bond<br>Donor (<5) | Hydrogen Bond<br>Acceptor (<10) | LogP<br>(<5) | Violations |
| --- | --- | --- | --- | --- | --- |
| **Vitamin E** | 430 | 1 | 2 | 8.840264 | 1 violation |
| **Nonacosane** | 408 | 0 | 0 | 9.128418 | 1 violation |
| **Stigmasterol** | 412 | 1 | 1 | 7.800803 | 1 violation |
| **BPA** | 292 | 1 | 3 | 4.092798 | 0 violations |

**BPA-** Benzenepropanoic acid, 3,5-bis(1,1-dimethylethyl)-4-hydroxy-, methyl ester

**Table 3.**
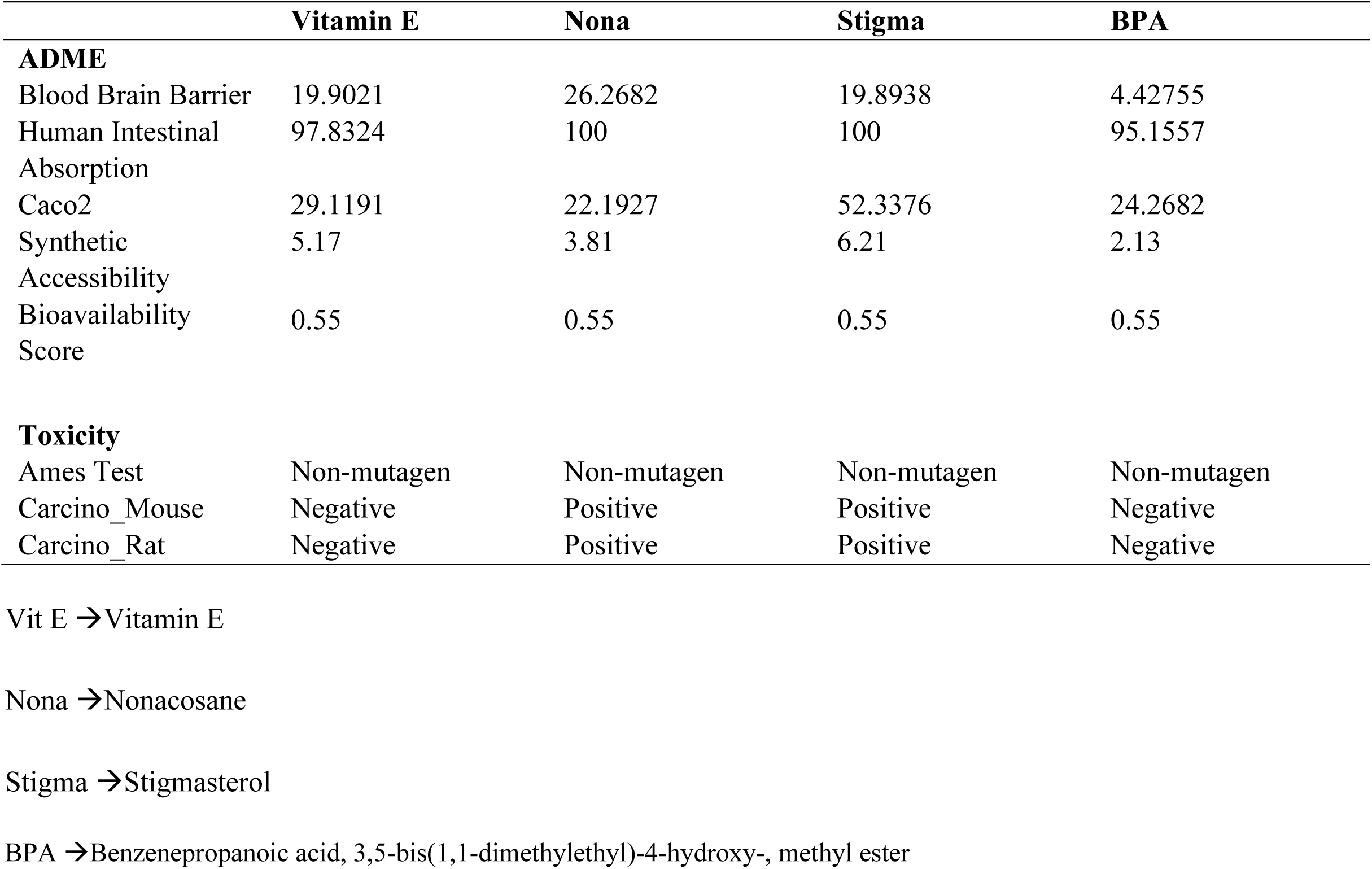
ADME results for the screened phytochemicals.

The lipophilicity (LogP) is an important factor to be considered while screening compounds based on their drug likeness ability. LogP values affect the rate of absorption of drug molecules in the body. The higher the value of LogP the lower absorption of the drug molecule. The effective range of logP value for anti-neurodegenerative drugs is in the range of 1-2 [44]. Although the logP values of the top hit phytochemicals were higher than the optimum values for anti-neurodegenerative drugs, an effective process of drug repurposing can be applied in order to bring down the logP value in the optimum range as some of the commercial drugs used against different neurodegenerative diseases shows the logP value of more than the optimum range (R-BPAP – 4.183, Valproic acid – 2.8106).

The Synthetic Accessibility (SA) value of a compound suggests the ease with which the compound can be synthesized. SA value of 1 suggests that the compound can be easily synthesized whereas the value of 10 suggests that the synthesis can be challenging. The SA values of the phytochemicals were in the range of 2.13-6.21. The Bioavailability Score (BS) represents the amount of drug that will reach the targeted site once the drug has been injected in the body. All the phytochemicals showed a BS of 0.55.

The Blood Brain Barrier (BBB) is an important factor to consider while developing a potential neurodegenerative drug. The BBB value of all the top hit phytochemicals falls in the category of CNSactive compound (Supplementary Information S2). Human Intestinal Absorption (HIA) value denotes the extent to which the drugs will be absorbed by the human system. All the top hit phytochemicals showed a very high HIA value suggesting a good absorption rate. Caco-2 values represents the ingestion of orally administered drugs. All the top hit phytochemicals represented a moderate permeability.

Toxicity is another important factor that has to be considered while synthesizing a new drug to target any disease. The most common type of toxicity tests are Ames test and Rodent Carcinogenicity. Ames test is a biological assay to examine the mutagenicity of a compound. All the top-hit phytochemicals are grouped in the non-mutagenic category. Carcinogenicity is a type of toxicity that induces cancer in the body. Vitamin E and BPA showed clear evidence of carcinogenic activity whereas, Stigmasterol and Nonacosane showed no evidence of carcinogenic activity.

## 4. DISCUSSION

Neurodevelopmental disorders are an important aspect to be considered. The disorders affect various parts of brain resulting in changes associated with memory, locomotion, behavior, emotions. Neurocognition and attention is an important aspect of humans to survive on a day-to-day basis. The binding behavior of neurotrophins with p75NTR receptor initiates a cascade of process that are fundamental for the neuronal signaling [45]. p75NTR interacts with tropomyosin related kinase (Trk) and modulate the Trk signaling but is also responsible for initiating the proapoptotic signaling process [46,47,48]. Thus, it is of utmost importance to devise drugs that can target the neurocognitive and neurodevelopmental disorders in order to assist the patient with a better lifestyle. Synthetic drugs possess the risk of side effect due to its chemically synthesized process. *Bacopa monnieri* has garnered much attention in recent years for its ability to enhance the neurocognitive and attention properties of human brain [45]. This study focuses on finding the phytochemicals of *B. monnieri* with potential to inhibit the activity of the causative protein in the case of Neurodevelopmental disorder. In the current investigation, we screened the 100 phytochemicals of *B. monnieri* and docked it against the Neurotrophins and identified the major phytochemicals potential to inhibit the activity of the Neurotrophins involved in the neurodegenerative diseases. The molecular docking studies illustrated that several phytochemicals of *B. monnieri* has a better binding affinity with Vitamin E, BPA, Stigmasterol and Nonacosane showing the best binding behavior with BDNF, NT3, NT4 and NGF respectively. The docking results (Fig. 2) showed the interaction of the respective phytochemicals with the active residues of the neurotrophins suggesting the potential of these phytochemicals in neurodegenerative disorders. Molecular docking analysis was followed by Molecular Dynamic (MD) Simulations for 150ns. MD Simulation studies revealed the stability of the system throughout the simulation run. Root Mean Square Deviation (RMSD) analysis showed that in the presence of the top-hit phytochemical, the Neurotrophins, BDNF, NT3 and NT4 remain stable throughout the MD run (Fig. 4a, 4b, 4c). NGF showed a fluctuation characteristic suggesting a lower binding with the ligand (Fig 4d). Root Mean Square Fluctuation (RMSF) analysis showed that the stability of the residues present in the active site remains stable when the top-hit phytochemicals bind to the Neurotrophins (Fig 5). Radius of Gyration (RoG) analysis further confirmed the stability of the Neurotrophins by showing a stable and compact progress throughout the run (Fig 6) followed by a constant Total Energy stability of the protein ligand system (Fig 7). Based on these study, four major phytochemicals were screened for ADME studies to find their characteristic likeness. All the four phytochemicals followed the major criteria of Drug likeness rule (Lipinski’s) by showing only 1 violation and in case of BPA no violation and the bioavailability score of these phytochemicals showed a good score suggesting their potential as a natural inhibitor (Table 1). The toxicity studies revealed less toxicity of the top hit phytochemicals with a major concern to be focused on the carcinogenicity activity of two phytochemicals (Table 1). Future work with animal models can provide us with experimental results that will further support the current study and take us a step closer for the development of neurodegenerative drugs.

## 5. CONCLUSION

*Bacopa monnieri* has been extensively used to treat the neurological disorders. Its ayurvedic roots make it a plant of medicinal purpose. It is associated with improving the brain function including intelligence and cognition [25]. *B. monnieri* extract when used to treat the animal model of Alzheimer’s (C57/Bl6 mice) showed a neuroprotective mechanism and it restricts the β-amyloid formation [46]. These results suggest that *B. monnieri* can act as a potential drug candidate against several other neurodegenerative disorders. Yet, it is unclear as to how *B. monnieri* actually causes these activities. The current work was done with aim of understanding the major involvement of phytochemical that proves *B. monnieri* to be useful in cases of neurodegenerative disorders. This study suggests that, BPA of *B. monnieri* possess potential to act as a drug candidate which can be employed for the treatment of Neurodevelopmental disorders. The binding behavior of Vitamin E, BPA, Stigmasterol and Nonacosane with the active residues of BDNF, NT3, NT4 and NGF respectively, shows the involvement of hydrophobic interactions along with positively charged interactions. These interactions reveal the microscopic scenario of how the phytochemicals interact with the active residues thus suggesting their potential role as inhibitors. Further molecular dynamic simulations suggested the stable binding behavior of the phytochemicals. Thus, laying down the foundation that these phytochemicals interact in a stable manner with the respective neurotrophins. Still, experiments will be required to find the molecular mechanism behind its action and to find out how the plant imparts its effect. Our work is a step towards unraveling the mystery behind the medicinal property of *B. monnieri*. The computational studies conducted in this work reveals the innate medicinal nature of the phytochemicals and their potential to act as a drug candidate.

## Abbreviations

BDNF: Brain-derived Neurotrophic Factor
NT3: Neurotrophin 3
NT4: Neurotrophin 4
NGF: Nerve Growth Factor
ND: Neurodegenerative disorders
BM: *Bacopa monnieri*
PDB: Protein Data Bank
RMSD: Root mean square deviation

## 7. ACKNOWLEDGEMENT

Authors acknowledge their respective affiliating institutions.

## 8. STATEMENTS & DECLARATIONS

### Ethical Approval

The present research study does not involve any human participants, their data or biological material.

### Consent to Participate

Not applicable

### Consent to Publish

Not applicable

### Authors Contributions

**SS, AK conceived the idea, SS, AK, did the experiments, SS, SM analyzed the results, SS, AK, SM prepared the manuscript, YKM, MS & NR added illustrations and figures, SS, HT, PKH reviewed the manuscript**.

### Availability of data and materials

Not Applicable

## REFERENCES

[1] B. Hempstead, Dissecting the diverse actions of pro- and mature neurotrophins, Curr. Alzheimer Res. 3 (2006) 19–24. https://doi.org/10.2174/156720506775697061.

[2] P.A. Leake, O. Akil, H. Lang, Neurotrophin gene therapy to promote survival of spiral ganglion neurons after deafness, Hear. Res. 394 (2020) 107955. https://doi.org/10.1016/J.HEARES.2020.107955.

[3] M.P. Sahu, Y. Pazos-Boubeta, C. Pajanoja, S. Rozov, P. Panula, E. Castrén, Neurotrophin receptor Ntrk2b function in the maintenance of dopamine and serotonin neurons in zebrafish, Sci. Reports 2019 91. 9 (2019) 1–13. https://doi.org/10.1038/s41598-019-39347-3.

[4] L.F. Reichardt, Neurotrophin-regulated signalling pathways, Philos. Trans. R. Soc. Lond. B. Biol. Sci. 361 (2006) 1545–1564. https://doi.org/10.1098/RSTB.2006.1894.

[5] A.E. Autry, D. Bambah-Mukku, The role of brain-derived neurotrophic factor in neural circuit development and function, Synap. Dev. Matur. (2020) 443–466. https://doi.org/10.1016/B978-0-12-823672-7.00020-X.

[6] M. Vilar, H. Mira, Regulation of Neurogenesis by Neurotrophins during Adulthood: Expected and Unexpected Roles, Front. Neurosci. 10 (2016). https://doi.org/10.3389/FNINS.2016.00026.

[7] A. Sahay, A. Kale, S. Joshi, Role of neurotrophins in pregnancy and offspring brain development, Neuropeptides. 83 (2020). https://doi.org/10.1016/J.NPEP.2020.102075.

[8] Konar, N. Shah, R. Singh, N. Saxena, S.C. Kaul, R. Wadhwa, M.K. Thakur, Protective role of Ashwagandha leaf extract and its component withanone on scopolamine-induced changes in the brain and brain-derived cells, PLoS One. 6 (2011). https://doi.org/10.1371/JOURNAL.PONE.0027265.

[9] P.T. Tseng, Y.W. Chen, K.Y. Tu, H.Y. Wang, W. Chung, C.K. Wu, S.P. Hsu, H.C. Kuo, P.Y. Lin, Statedependent increase in the levels of neurotrophin-3 and neurotrophin-4/5 in patients with bipolar disorder: A meta-analysis, J. Psychiatr. Res. 79 (2016) 86–92. https://doi.org/10.1016/J.JPSYCHIRES.2016.05.009.

[10] Y.C. Kim, Neuroprotective phenolics in medicinal plants, Arch. Pharm. Res. 33 (2010) 1611–1632. https://doi.org/10.1007/S12272-010-1011-X

[11] V.G. Gómez-Pineda, F.M. Torres-Cruz, C.I. Vivar-Cortés, E. Hernández-Echeagaray, Neurotrophin-3 restores synaptic plasticity in the striatum of a mouse model of Huntington’s disease, CNS Neurosci. Ther. 24 (2018) 353–363. https://doi.org/10.1111/CNS.12824.

[12] S.J. Allen, D. Dawbarn, Clinical relevance of the neurotrophins and their receptors, Clin. Sci. (Lond). 110 (2006) 175–191. https://doi.org/10.1042/CS20050161.

[13] T. Cho, J.K. Ryu, C. Taghibiglou, Y. Ge, A.W. Chan, L. Liu, J. Lu, J.G. McLarnon, Y.T. Wang, Long-term potentiation promotes proliferation/survival and neuronal differentiation of neural stem/progenitor cells, PLoS One. 8 (2013). https://doi.org/10.1371/JOURNAL.PONE.0076860.

[14] C.P. Fitzsimons, E. Van Bodegraven, M. Schouten, R. Lardenoije, K. Kompotis, G. Kenis, M. Van Den Hurk, M.P. Boks, C. Biojone, S. Joca, H.W.M. Steinbusch, K. Lunnon, D.F. Mastroeni, J. Mill, P.J. Lucassen, P.D. Coleman, D.L.A. Van Den Hove, B.P.F. Rutten, Epigenetic regulation of adult neural stem cells: implications for Alzheimer’s disease, Mol. Neurodegener. 9 (2014). https://doi.org/10.1186/17501326-9-25.

[15] D.A. Simmons, Modulating Neurotrophin Receptor Signaling as a Therapeutic Strategy for Huntington’s Disease, J. Huntingtons. Dis. 6 (2017) 303–325. https://doi.org/10.3233/JHD-170275.

[16] J. Meldolesi, Neurotrophin receptors in the pathogenesis, diagnosis and therapy of neurodegenerative diseases, Pharmacol. Res. 121 (2017) 129–137. https://doi.org/10.1016/J.PHRS.2017.04.024.

[17] Y. Jiang, J.M. Fay, C.D. Poon, N. Vinod, Y. Zhao, K. Bullock, S. Qin, D.S. Manickam, X. Yi, W.A. Banks, A. V. Kabanov, Nanoformulation of Brain-Derived Neurotrophic Factor with Target Receptor-Triggered-Release in the Central Nervous System, Adv. Funct. Mater. 28 (2018). https://doi.org/10.1002/ADFM.201703982.

[18] I.L. Shamovsky, D.F. Weaver, G.M. Ross, R.J. Riopelle, The interaction of neurotrophins with the p75NTR common neurotrophin receptor: a comprehensive molecular modeling study, Protein Sci. 8 (1999) 2223–2233. https://doi.org/10.1110/PS.8.11.2223.

[19] P. Kuner, C. Hertel, NGF Induces Apoptosis ina Human Neuroblastoma Cell Line Expressing the Neurotrophin Receptor p75NTR., J NeuorSci Res. 54(1998) 465–74. https://doi.org/10.1002/(SICI)1097-4547(19981115)54:4

[20] A. Colquhoun, G.M. Lawrance, I.L. Shamovsky, R.J. Riopelle, G.M. Ross, Differential activity of the nerve growth factor (NGF) antagonist PD90780 [7-(benzolylamino)-4,9-dihydro-4-methyl-9-oxo-pyrazolo[5,1b]quinazoline-2-carboxylic acid] suggests altered NGF-p75NTR interactions in the presence of TrkA, J. Pharmacol. Exp. Ther. 310 (2004) 505–511. https://doi.org/10.1124/JPET.104.066225.

[21] M.K. Saraf, S. Prabhakar, A. Anand, Neuroprotective effect of Bacopa monniera on ischemia induced brain injury, Pharmacol. Biochem. Behav. 97 (2010) 192–197. https://doi.org/10.1016/J.PBB.2010.07.017.

[22] C. Stough, A. Scholey, V. Cropley, K. Wesnes, A. Zangara, M. Pase, K. Savage, K. Nolidin, J. Lomas, L.A. Downey, Examining the cognitive effects of a special extract of Bacopa monniera (CDRI 08: KeenMind): A review of ten years of research at Swinburne University, J. Pharm. Pharm. Sci. 16 (2013) 254–258. https://doi.org/10.18433/J35G6M.

[23] S.K. Bhattacharya, S. Ghosal, Anxiolytic activity of a standardized extract of Bacopa monniera: an experimental study., Phytomedicine. 5 (1998) 77–82. https://doi.org/10.1016/S0944-7113(98)80001-9.

[24] G.K. Shinomol, R.B. Mythri, M.M. Srinivas Bharath, Muralidhara, Bacopa monnieri extract offsets rotenone-induced cytotoxicity in dopaminergic cells and oxidative impairments in mice brain, Cell. Mol. Neurobiol. 32 (2012) 455–465. https://doi.org/10.1007/S10571-011-9776-0.

[25] N.P. Sukumaran, A. Amalraj, S. Gopi, Neuropharmacological and cognitive effects of Bacopa monnieri (L.) Wettst - A review on its mechanistic aspects, Complement. Ther. Med. 44 (2019) 68–82. https://doi.org/10.1016/J.CTIM.2019.03.016.

[26] A. Russo, F. Borrelli, Bacopa monniera, a reputed nootropic plant: an overview, Phytomedicine. 12 (2005) 305–317. https://doi.org/10.1016/J.PHYMED.2003.12.008.

[27] S. Saravana Kumar, P. Saraswathi, R. Vijayaraghavan, Effect of Bacopa Monniera on Cold Stress Induced Neurodegeneration in Hippocampus of Wistar Rats: A Histomorphometric Study, J. Clin. Diagn. Res. 9 (2015) AF05. https://doi.org/10.7860/JCDR/2015/10199.5423.

[28] P. Jain, H.P. Sharma, F. Basri, K. Priya, P. Singh, Phytochemical analysis of Bacopa monnieri (L.) Wettst. and their anti-fungal activities, Indian J. Tradit. Knowl. 16 (2017) 310–318.

[29] S. Kim, J. Chen, T. Cheng, A. Gindulyte, J. He, S. He, Q. Li, B.A. Shoemaker, P.A. Thiessen, B. Yu, L. Zaslavsky, J. Zhang, E.E. Bolton, PubChem 2019 update: improved access to chemical data, Nucleic Acids Res. 47 (2019) D1102–D1109. https://doi.org/10.1093/NAR/GKY1033.

[30] N.M. O’Boyle, M. Banck, C.A. James, C. Morley, T. Vandermeersch, G.R. Hutchison, Open Babel: An Open chemical toolbox, J. Cheminform. 3 (2011). https://doi.org/10.1186/1758-2946-3-33.

[31] ChemOffice, 7.0.1. 2002. CambridgeSoft, Corporation, Cambridge, MA.

[32] E.F. Pettersen, T.D. Goddard, C.C. Huang, G.S. Couch, D.M. Greenblatt, E.C. Meng, T.E. Ferrin, UCSF Chimera--a visualization system for exploratory research and analysis, J. Comput. Chem. 25 (2004) 1605–1612. https://doi.org/10.1002/JCC.20084.

[33] Schrodinger, LLC 2010. The PyMOL Molecular Graphics System, Version 1.8.4.0

[34] Duhovny, R. Nussinov, H.J. Wolfson, Efficient unbound docking of rigid molecules, Lect. Notes Comput. Sci. (Including Subser. Lect. Notes Artif. Intell. Lect. Notes Bioinformatics). 2452 (2002) 185–200. https://doi.org/10.1007/3-540-45784-4_14.

[35] Schneidman-Duhovny, Y. Inbar, R. Nussinov, H.J. Wolfson, PatchDock and SymmDock: servers for rigid and symmetric docking, Nucleic Acids Res. 33 (2005). https://doi.org/10.1093/NAR/GKI481.

[36] M.J. Abraham, T. Murtola, R. Schulz, S. Páll, J.C. Smith, B. Hess, E. Lindah, GROMACS: High performance molecular simulations through multi-level parallelism from laptops to supercomputers, SoftwareX. 1–2 (2015) 19–25. https://doi.org/10.1016/J.SOFTX.2015.06.001.

[37] B. Hess, C. Kutzner, D. Van Der Spoel, E. Lindahl, GRGMACS 4: Algorithms for highly efficient, loadbalanced, and scalable molecular simulation, J. Chem. Theory Comput. 4 (2008) 435–447. https://doi.org/10.1021/CT700301Q.

[38] Van Der Spoel, E. Lindahl, B. Hess, G. Groenhof, A.E. Mark, H.J.C. Berendsen, GROMACS: fast, flexible, and free, J. Comput. Chem. 26 (2005) 1701–1718. https://doi.org/10.1002/JCC.20291.

[39] S.K. Das, S. Mahanta, B. Tanti, H. Tag, P.K. Hui, Identification of phytocompounds from Houttuynia cordata Thunb. as potential inhibitors for SARS-CoV-2 replication proteins through GC-MS/LC-MS characterization, molecular docking and molecular dynamics simulation, Mol. Divers. (2021). https://doi.org/10.1007/S11030-021-10226-2.

[40] J.D. Hunter, Matplotlib, Comput. Sci. Eng. 9 (2007) 90–95. https://doi.org/10.1109/MCSE.2007.55.

[41] C.A. Lipinski, Lead- and drug-like compounds: the rule-of-five revolution, Drug Discov. Today Technol. 1 (2004) 337–341. https://doi.org/10.1016/J.DDTEC.2004.11.007.

[42] A. Daina, O. Michielin, V. Zoete, SwissADME: a free web tool to evaluate pharmacokinetics, drug-likeness and medicinal chemistry friendliness of small molecules, Sci. Reports 2017 71. 7 (2017) 1–13. https://doi.org/10.1038/srep42717.

[43] S. K. Lee, I. H. Lee, H. J. Kim, G. S. Chang, J. E. Chung, K. T. No, The PreADME Approach: Web Based program for rapid prediction of physico-chemical, drug absorption and drug-like properties. EuroQSAR 2002 Designing Drugs and Crop Protectants: processes, problem and solutions (2003) 418–420. Blackwell Publishing, Massachusetts, USA

[44] H. Pajouhesh, G.R. Lenz, Medicinal chemical properties of successful central nervous system drugs, NeuroRx. 2 (2005) 541–553. https://doi.org/10.1602/NEURORX.2.4.541.

[45] E.J. Huang, L.F. Reichardt, Neurotrophins: roles in neuronal development and function, Annu. Rev. Neurosci. 24 (2001) 677–736. https://doi.org/10.1146/annurev.neuro.24.1.677

[46] A.H. Salehi, P.P. Roux, C.J. Kubu, C. Zeindler, A. Bhakar, L.L. Tannis, J.M Verdi, P.A. Barker, NRAGE, a novel MAGE protein, interacts with the p75 neurotrophin receptor and facilitates nerve growth factordependent apoptosis, Neuron 27 (2000) 279–288. https://doi.org/10.1038/cdd.2008.127

[47] P.P. Roux, A.L. Bhakar, T.E. Kennedy, P.A. Barker, The p75 neurotrophin receptor activates Akt (protein kinase B) through a phosphatidylinositol 3-kinase-dependent pathway, J Biol Chem. 276(25) (2001) 23097–104. https://doi.org/10.1074/jbc.M011520200

[48] V. Mamidipudi, x, Li, M.W. Wooten, Identification of interleukin 1 receptor-associated kinase as a conserved component in the p75-neurotrophin receptor activation of nuclear factor-kappa B. J Biol Chem. 277(31) (2002) 28010–8. https://doi.org/10.1074/jbc.M109730200

[49] K.S. Chaudhari, N.R. Tiwari, R.R. Tiwari, R.S. Sharma, Neurocognitive effect of nootropic drug Brahmi (Bacopa monnieri) in Alzheimer’s disease, Ann. Neurosci. 24 (2017) 111–122. https://doi.org/10.1159/000475900.

[50] M. Dhanasekaran, B. Tharakan, L.A. Holcomb, A.R. Hitt, K.A. Young, B. V. Manyam, Neuroprotective mechanisms of ayurvedic antidementia botanical Bacopa monniera, Phyther. Res. 21 (2007) 965–969. https://doi.org/10.1002/PTR.2195.

