## Supplementary Information for "*In silico* analysis of *Bacopa monnieri* (L.) Wettst. compounds for drug development against Neurodegenerative Disorders"

**This file includes**

Supporting Information:

- Detailed Methodology for Molecular Docking
- Detailed methodology for ADME-Tox studies
- Molecular Docking Analysis of other prominent phytochemicals
- Table S6 – Molecular Docking interactions of other prominent phytochemicals with respective Neurotrophins
- Table S7 – Molecular weight and Structure of 22 phytochemicals apart from the top hit phytochemicals
- Figure S1 – Crystal structure of the Neurotrophins
- Free Energy Surface Plot of Neurotrophins
- Figure S2 – Free energy surface plot of Neurotrophins with corresponding structure of ligand

### S1. Detailed Methodology for Molecular Docking

*S1.1. Energy Minimization using MM2*

MM energy minimization stands for Molecular Mechanics energy minimization. Force field calculations provides the choice for the molecular structure and energy determination under various circumstances. MM2 energy minimization is an upgraded form of MM1 energy minimization proposed by Norman L. Allinger in 1977 (Allinger, 1977). MM2 energy minimization was developed in order to undertake the minimization of the hydrocarbons. The difference between MM1 and MM2 energy minimization includes a different writing terminology for the torsional energy represented as:

$$E\omega = \frac{V1}{2}(1 + cos\omega) + \frac{V2}{2}(1 + cos2\omega) + \frac{V3}{2}(1 + cos3\omega)$$

Where ω is always measured from 0° to 180°.

*S1.2. Docking Study*

PatchDock server (bioinfo3d.cs.tau.ac.il/PatchDock/php.php) (Duhovny et al., 2002; SchneidmanDuhovny et al., 2005) was utilized for the molecular docking studies. PatchDock server requires two input: the input molecule can be of any type either protein, DNA, peptide or drug. The output generated is an ensemble of potential complexes that are sorted based on the shape complementarity principle. When two molecules are supplied to the PatchDock server it initiates a process of three steps.

Step 1: Molecular Shape Representation

In this step molecular surface topology is computed in order to assess the nature of molecular surface. A segmentation algorithm detects that convex, concave and flat surfaces. The patches are then filtered in order to retain the hot-spot residues.

Step 2: Surface Patch Matching

Geometric Hashing and Pose-Clustering matching techniques are applied to match the patches, Concave with convex and flat surfaces with any surface.

Step 3: Filtering and Scoring

The unacceptable structures are discarded and only the relevant structures are listed out. The output generated by the PatchDock is analyzed using two types of scores. PatchDock score gives the geometric scores based on the surface complementarity of the molecule. The higher the PatchDock score the better the shape complementarity. ACE (Atomic Contact Energy) is the desolvation free energy required to transfer atoms from water to protein’s interior.

### S2. Detailed Methodology for ADME-Tox analysis

Adsorption, Digestion, Metabolism, Excretion and Toxicity provides a detailed analysis of pharmacokinetic characteristics of a compound posing a potential to act as a drug molecule. The ADME-Tox study was performed using the PreADMET server (https://preadmet.bmdrc.kr/). The PreADMET server provides several characteristics of the compound along with a detailed summary of how to interpret the results.

S2.1. Blood-Brain Barrier:

Blood Brain Barrier (BBB) values represent whether a compound has the potential to penetrate the barrier and effectively show its activity. Usually BBB values are calculated for the drugs that are targeted for the treatment of Neurodegenrative disorders so that they can effectively penetrate the Central Nervous System (CNS). BBB values are calculated as the ratio of the concentration of the radiolabelled compounds in brain and blood (Ajay et al., 1999).

$$BBB = \frac{[Brain]}{[Blood]}$$

PreADMET server classifies the molecule as either CNS-active compound or CNS-inactive compound. Compounds classified as CNS-active must pass the BB barrier whereas the CNS-inactive compound mustn't cross the BB barrier. PreADMET classification is as follows:

Table S1: Classification of compounds for Blood-Brain Barrier according to PreADMET

| Classification | BB Value |
| --- | --- |
| High Absorption to CNS | More than 2.0 |
| Middle Absorption to CNS | 2.0 ~ 0.1 |
| Low Absorption to CNS | Less than 0.1 |

S2.2. Human Intestinal Absorption:

Human Intestinal Absorption (HIA) is another important factor to consider while screening compounds for drug discovery. HIA values suggest the ability of the compound to get effectively absorbed from the human gastrointestinal system into the bloodstream. PreADMET calculates the HIA of the compound at pH 7.4 as the HIA's are measured via *in vivo* test (Zhao et al., 2001). PreADMET classification for HIA is as follows:

Table S2: Classification of compounds for Human Intestinal Absorption according to PreADMET

| Classification | Human Intestinal Absorption (HIA) |
| --- | --- |
| Poorly Absorbed Compounds | 0 ~20 % |
| Moderately Absorbed Compounds | 20 ~ 70 % |
| Well Absorbed Compounds | 70 ~ 100 % |

S2.3. Caco2:

Caco2 is a defined cell line of human colorectal adenocarcinoma cells. It is used as a model of intestinal epithelial barrier. Caco2 cells are used as a monolayer on a cell culture and is broadly utilized in the pharmaceutical industries as an in vitro model of human small digestive tract mucosa to foresee the ingestion of orally regulated medications. PreADMET server predicts the Caco-2 cell permeability by utilizing the chemical structure if the compounds at pH 7.4 as the permeability of Caco-2 cells are measured at this pH (Yamashita et al., 2000). PreADMET classification for Caco-2 cell permeability is as follows:

Table S3: Classification of compounds for Caco-2 according to PreADMET

| Classification | $P_{Caco-2}$ (nm/sec) |
| --- | --- |
| Low permeability | Less than 4 |
| Moderate permeability | 4 ~ 70 |
| High Permeability | More than 70 |

S2.4. Ames Test:

Ames test is a biological assay to examine the mutagenicity of a chemical compound. Ames test is performed by utilizing several strains of *Salmonella typhimurium* carrying mutations in those genes that are involved in the Histidine synthesis. Thus, the strains used are "auxotrophic" i.e., they require Histidine for growth but cannot produce it. This method tests the capability of the chemical compound to bring about a mutation such that the selected strains can now grow in a Histidine free media. A positive Ames test indicates that the chemical compound being tested is mutagenic and can thus be characterized as a carcinogen (Ames et al., 1972). The PreADMET server classifies the Ames test as:

Table S4: Classification of compounds for Ames Test according to PreADMET

| Classification | Test Type |
|---|---|
| Non-mutagen | Negative |
| Mutagen | Positive |

S2.5. Rodent Carcinogenicity:

Carcinogenicity is a type of toxicity that induces cancer in the body. For a long time people have been using the "2-year carcinogenicity test" to predict whether a compound is a potential carcinogen or not. Generally these studies are done on rodents to observe the existence of cancer. PreADMET predicts the results from its model that is based on the data of NTP (National Toxicology Program) and US FDA which comprises of in vivo results on mice and rats for 2 years. PreADMET classifies that results in following way:

Table S5: Classification of compounds for Rodent Carcinogenicity according to PreADMET

| Classification | NTP Definition |
|---|---|
| Negative | Clear evidence of carcinogenic activity |
| Positive | No evidence of carcinogenic activity |

**S3. Docking Analysis of other prominent phytochemicals**

The residue interaction with other phytochemicals and their respective Neurotrophins are shown in Table S1. Apart from the top hit phytochemicals, whose details are presented in the main file, there were other major phytochemicals which showed an active interaction with their respective Neurotrophins. 3,7,11,15-tetramethyl-2-Hexadecen-1-ol, the second-best phytochemical, showed a notable interaction with BDNF by forming a hydrogen bond interaction with Arg88 and hydrophobic interactions with Trp19, Leu49 and Tyr52 with a docking score of 4,108 and an ACE value of -192.86 kcal/mol.

Similarly, 3a(1H)-Azulenol-2,3,4,5,8,8a-hexahydro-6,8a-dimethyl-3-(1-M showed an eminent interaction with NT3 by forming hydrogen bond with Gln83 and Arg103. It also showed hydrophobic interactions with Ala27 and Ile104 with a docking score of 2,890 and an ACE value of -36.87 kcal/mol.

2,6,10-Trimethyl,14-ethylene-14-Pentadecne showed interaction with NT4 by forming hydrophobic interactions with Val44, Ala47, Arg98, Leu52 and Trp110 with a docking score of 5,550 and an ACE of -213.87 kcal/mol. 3,7,11,15-tetramethyl-2-Hexadecen-1- ol showed carbon hydrogen bonding with Gln54 and hydrophobic interactions with Arg98, Ala47, Val97, Val44, Trp110, Trp112 and Phe56 with a docking score of 5,432 and an ACE value of -286.35 kcal/mol. 9-Octadecenoic acid (Z)- showed hydrogen bonding with Trp110 and hydrophobic interactions with Arg98, Ala47, Val97, Trp110 and Phe56 with a docking score of 5,398 and an ACE of -265.53 kcal/mol. Phytol showed hydrophobic interactions with Val44, Arg98, Ala47, Val97, Phe56 and Trp110 with a docking score of 5,550 and an ACE of -312.72 kcal/mol.

Heneicosane showed a vital interaction with NGF in the form of hydrophobic interactions with Met37, Leu39, Met92 and Lys95 with a docking score of 5,136 and an ACE of -147.24 kcal/mol.

**Table S6: Molecular docking of phytocompounds from *B. monnieri* against the selected receptors**

| Neurotrophins | Phytochemicals | PatchDock Score | ACE (kcal/mol) | Interacting Residues |
|---|---|---|---|---|
| **BDNF** | 3,7,11,15-tetramethyl-2Hexadecen-1-ol | 4108 | -192.86 | Trp19, Tyr52, Arg88, Leu49 |
| **NT3** | 3A(1H)-Azulenol,2,3,4,5,8,8Ahexahydro-6,8Adimethyl-3-(1M | 2890 | -36.87 | Gln83, Arg103, Ala27, Ile104 |
| **NT4** | 1,2-Benzene dicarboxylic acid, mono (2-ethylhexyl) ester | 4636 | -201.71 | Ala47, Trp110, Val44, Leu52, Alr98, Trp112, Phe56 |
| **NT4** | 2-Cyclohexen-1one, 4hydroxy-3,5,5-trimethyl- 4-(3oxo-1-butenyl)- | 4150 | -203.02 | Arg98, Trp110, Leu100, Gln54, Trp112 |
| **NT4** | 2-Cyclohexen-1-one, 3-(3-hydroxybutyl)-2,4,4trimethyl- | 3974 | -175.74 | Phe56, Trp112, Arg98, Trp110, Val97, Tyr55 |

| | | | | |
|---|---|---|---|---|
| **NT4** | 2-Nonenal, 2-Pentyl- | 4566 | -188.57 | Phe56, Arg98, Trp110, Val44 |
| **NT4** | 2-Pentadecanone, 6,10,14-Trimethyl- | 4890 | -200.07 | Arg98, Trp110, Lue52, Val44, Ala47 |
| **NT4** | 3,7,11,15-tetramethyl2-Hexadecen-1-ol | 5432 | -286.35 | Phe56, Val97, Trp112, Arg98, Trp110, Gln54, Ala47, Val44 |
| **NT4** | 3A(1H)-Azulenol,2,3,4,5,8,8 Ahexahydro-6,8Adimethyl-3-(1-M | 4016 | -215.66 | Val97, Arg98, Trp110, Val44 |
| **NT4** | 9-Octadecenoic acid (Z)- | 5398 | -265.53 | Val97, Phe56, Arg98, Trp110, Ala47 |
| **NT4** | Benzenepropanoic acid,3,5-bis(1,1dimethylethyl)-4hydroxy-, methyl ester | 4746 | -228.88 | Leu52, Trp110, Arg98 |
| **NT4** | Cis-9-Hexadecenal | 5076 | -183.13 | Arg98, Trp110, Leu52, Val44 |
| **NT4** | Cis-10-Nonadecenoic acid | 4750 | -294.98 | Phe56, Val97, Arg98, Trp112, Val44, Trp110, Ala47, Ala46, Pro45, Val44 |
| **NT4** | Dodecane | 4098 | -179.79 | Trp110, Val97, Phe56, Val44, Ala47, Arg98 |
| **NT4** | Phenol, 2-methoxy-4-(2-Propenyl)- | 3730 | -137.88 | Tyr55, Arg98, Trp110 |

| | | | | |
|---|---|---|---|---|
| **NT4** | Tridecane | 4432 | -173.83 | Trp110, Ala47, Val44, Phe56, Arg98, Trp112 |
| **NGF** | 2-Nonenal, 2-Pentyl- | 3996 | -126.33 | His84, Thr83, Val109, Ser17 |
| **NGF** | 2-Pentadecanone, 6,10,14-Trimethyl- | 4056 | -25.16 | Phe49, Ala98, Arg100, Phe101, Trp21, Gln96 |
| **NGF** | Cis-9-Hexadecenal | 3930 | -104.84 | Phe49, Ala98, Arg100, Phe101, Ile31, Trp21 |
| **NGF** | Hexadecanoic acid, 2hydroxy-1-(hydroxymethyl) ethyl ester | 4734 | -93.22 | Phe101, Arg100, Ala98, Gln96 |
| **NGF** | Hexadecanoic acid, methyl ester | 4372 | -64.23 | Phe49, Phe101, Arg100, Ala98, Gln96 |
| **NGF** | Octadecanoic acid | 4438 | -145.41 | Gly40, Leu39, Met37, Met92, Lys95 |
| **NGF** | Tridecane | 4782 | -337.63 | Phe101, Arg100, Ala98, Gln96 |
| **NGF** | Vitamin E | 4636 | -203.65 | Arg103, His84, Phe86, Val109, Val111, Phe54, Ser19, Ser17 |

**Table S7: The following table shows the Molecular weight and the corresponding chemical structure of other prominent phytochemicals.**

| Phytochemical Name | Molecular Weight (g/mol) | Phytochemical Structure |
|---|---|---|
| 1,2-Benzenedicarboxylic acid, mono(2-ethylhexyl) ester | 278.34 | 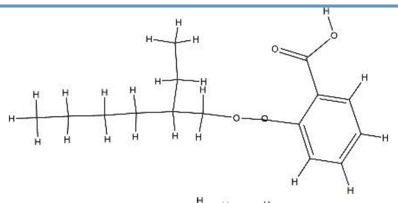 |
| 2,6,10-Trimethyl,14-ethylene-14-Pentadecne | | 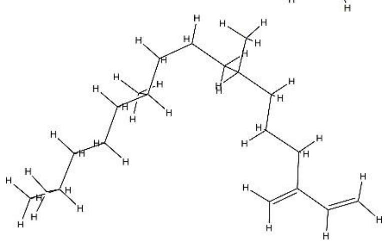 |
| 2-Cyclohexen-1-one,3-(3-hydroxybutyl)-2,4,4trimethyl- | 210.31 | 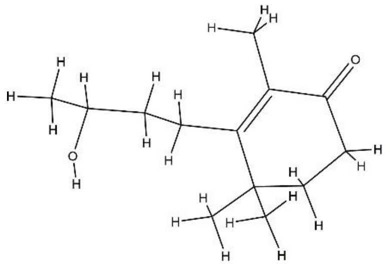 |
| 2-Cyclohexen-1-one, 4-hydroxy-3,5,5-trimethyl-4-(3oxo-1-butenyl)- | 222.28 | 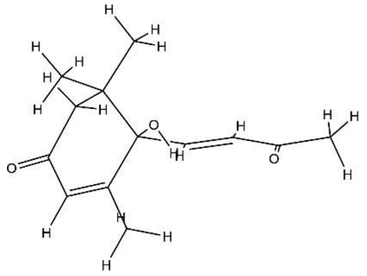 |
| 2-Nonenal, 2-Pentyl- | 210.36 | 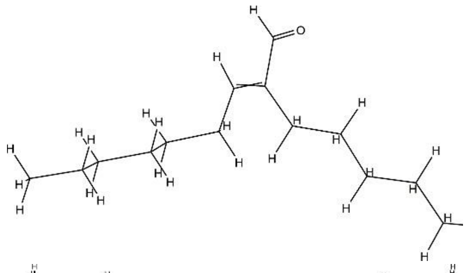 |
| 2-Pentadecanone,6,10,14-Trimethyl- | 268.47 | 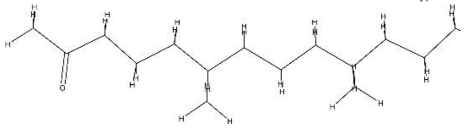 |

| | | |
|---|---|---|
| 3,7,11,15-tetramethyl-2-Hexadecen-1-ol | 296.5 | 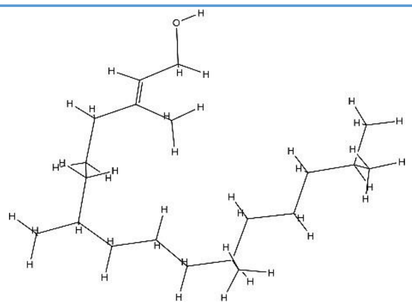 |
| 3a(1H)-Azulenol-2,3,4,5,8,8a-hexahydro-6,8a-dimethyl-3-(1-M | 222.37 | 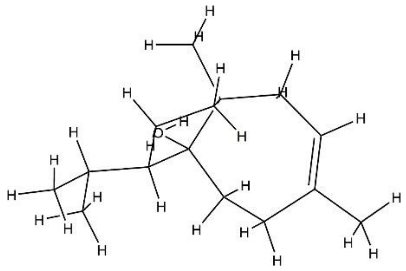 |
| 9-Octadecenoic acid (Z)- | 564.9 | 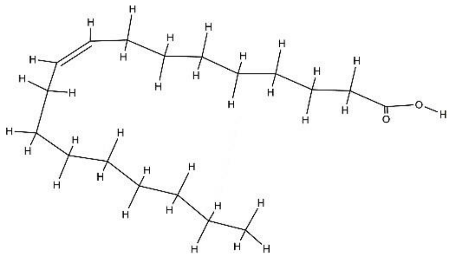 |
| 17-(1,5-Dimethyl-hex-2-enyl)- 10,13-dimethyl- 2,3,4,9,10,11,12,13 | 382.62 | 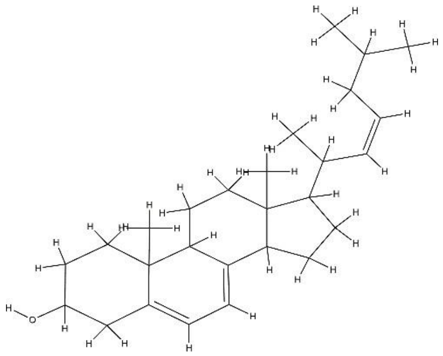 |
| Cis-9-Hexadecenal | 238.41 | 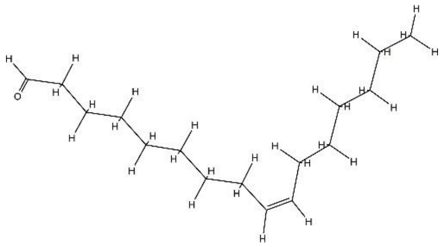 |

| | | |
|---|---|---|
| Cis-10-Nonadecenoic acid | 296.5 | 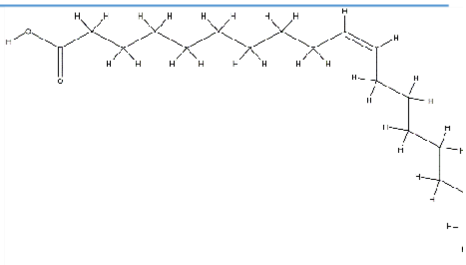 |
| Dodecane | 170.33 | 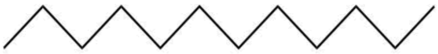 |
| Heneicosane | 296.6 | 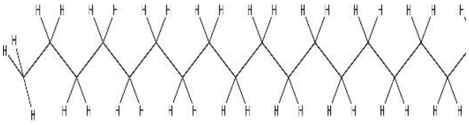 |
| Hexadecanoic acid, 2-hydroxy-1-(hydroxymethyl)ethyl ester | 330.5026 | 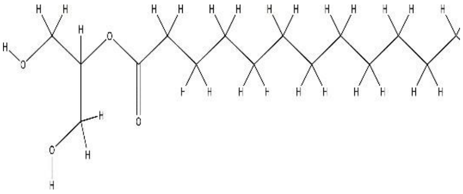 |
| Hexadecanoic acid, methyl ester | 270.4507 | 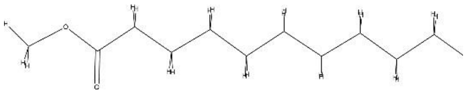 |
| Icosanoic acid | 312.5 | 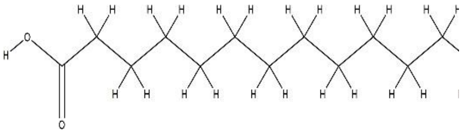 |
| Octadecanoic acid | 284.5 | 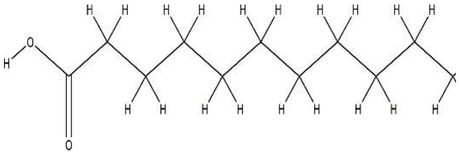 |
| Octadecanoic acid, ethyl ester | 312.5304 | 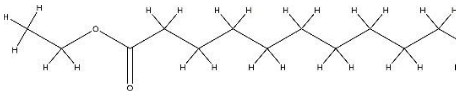 |
| Phenol, 2-methoxy-4-(2-Propenyl) | 162.4 | 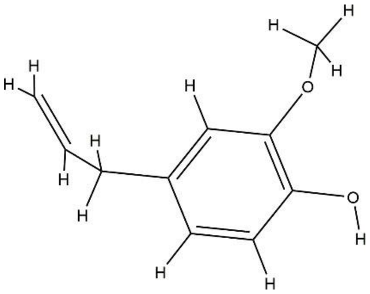 |

| | | |
|---|---|---|
| Phytol | 296.5 | 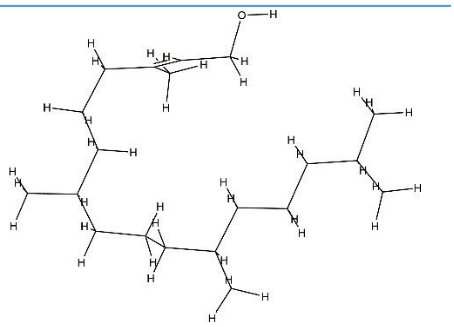 |
| Tridecane | 184.36 | 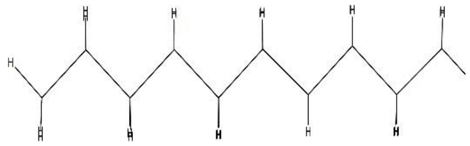 |

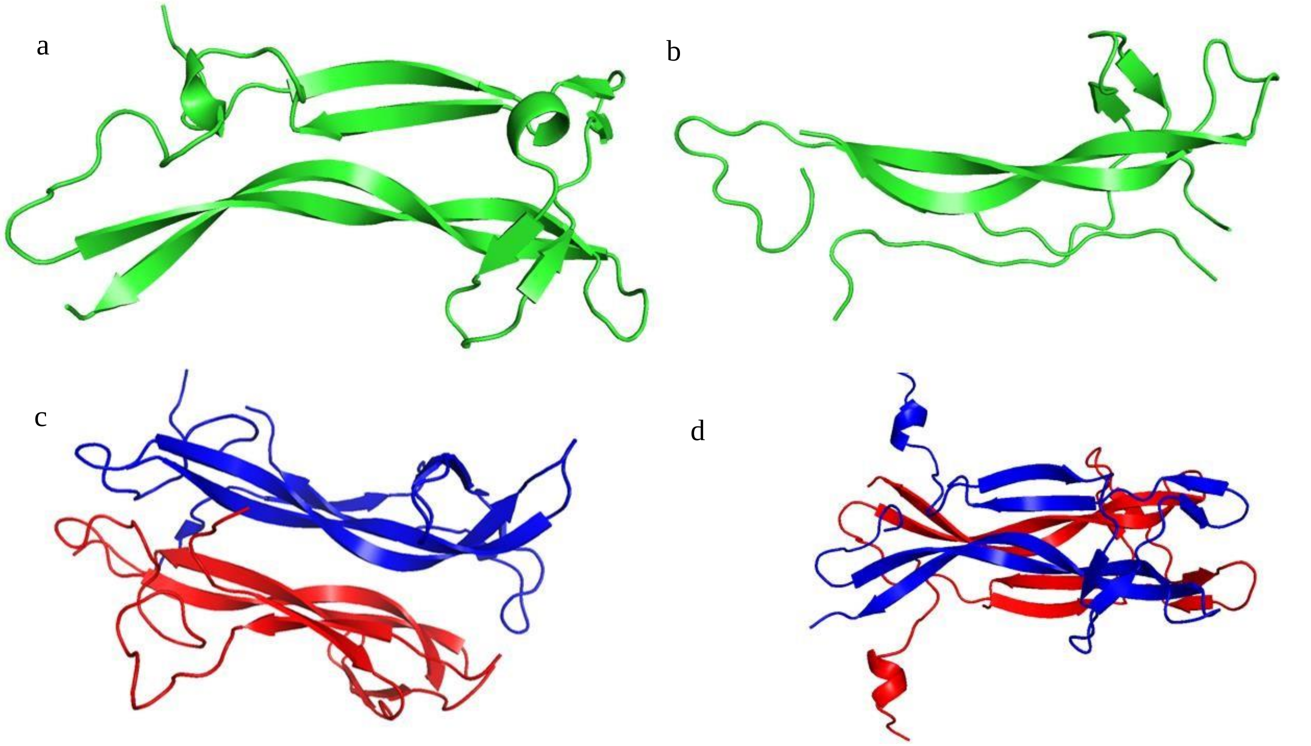


Figure S1: Crystal structure of receptors (a) BDNF (b) NT3 (c) NT4 (d) NGF. BDNF and NT3 consists of single chain represented in green color. NT4 and NGF are Homodimer where in the case of NT4 Chain A and Chain M are represented in red and blue color respectively. In the case of NGF Chain V and Chain W are represented in red and blue color respectively.

### S4. Free Energy Surface Plot

Figure S2 represents the Free Energy Surface (FES) diagram for the respective Neurotrophins. The FES of the respective Neurotrophins denote the dihedral angles plotted with respect to the free energy change. X and Y axis corresponds to the phi and psi angles and the corresponding color bar represents the free energy change. The most stable structural conformation of the system for all the Neurotrophins falls in the range of -140° to -150°Φ angle and 100° to 150°Ψ angle. In case of BDNF (Fig.S2a) the structural conformation of Vitamin E assumes the shape corresponding to nearly 140ns suggesting a stable interaction and the population of this conformation dominating the stable region. The structural integrity of BPA assumes that of the most stable structure at 140ns (Fig.S2b). This is evident as the binding stability of NT3 and BPA (Fig.4b) remained stabilized throughout the simulation time. Structure of Stigmasterol corresponding to 50ns dominated the region of most stabilized free energy. The end tail fluctuation of Stigmasterol might correspond to the regions showing faded purple color (Fig.S2c). Nonacosane assumed the structure (Fig.S2d) almost similar to the one corresponding to 30ns (Fig.3d). This is evident as the corresponding RMSD fluctuation of Nonacosane corresponding to 10-30ns duration is the most stable one.

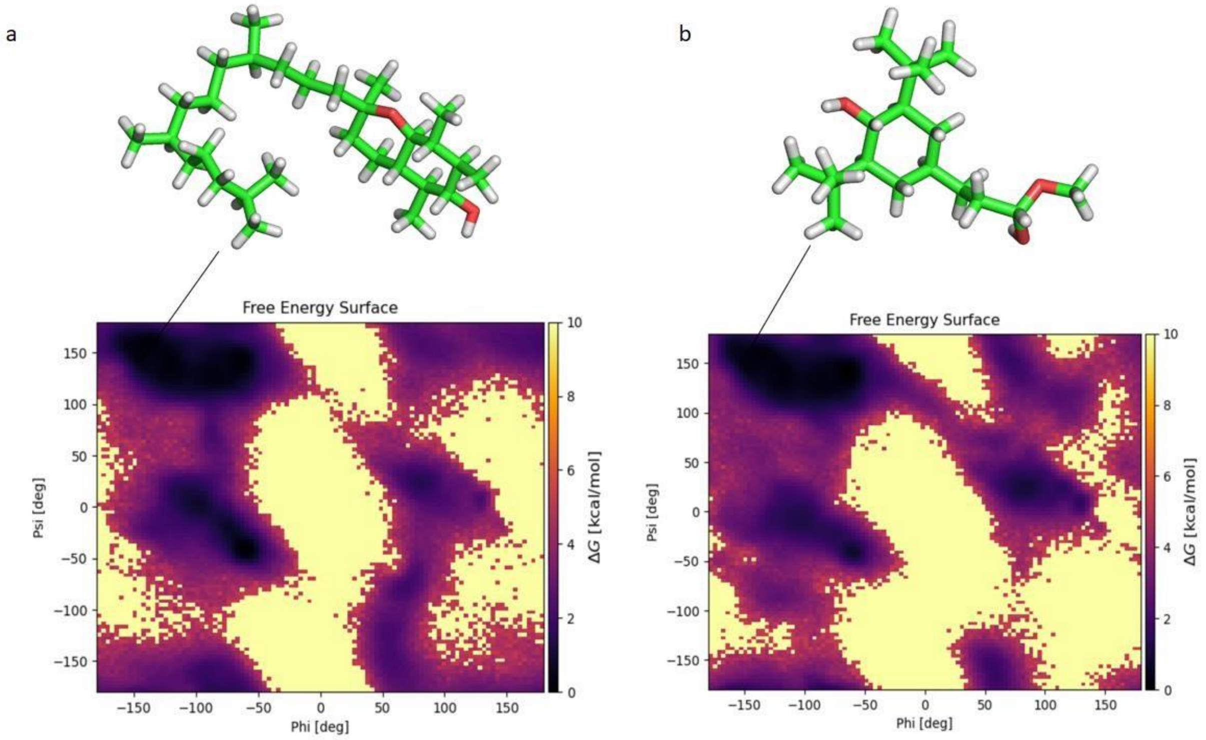

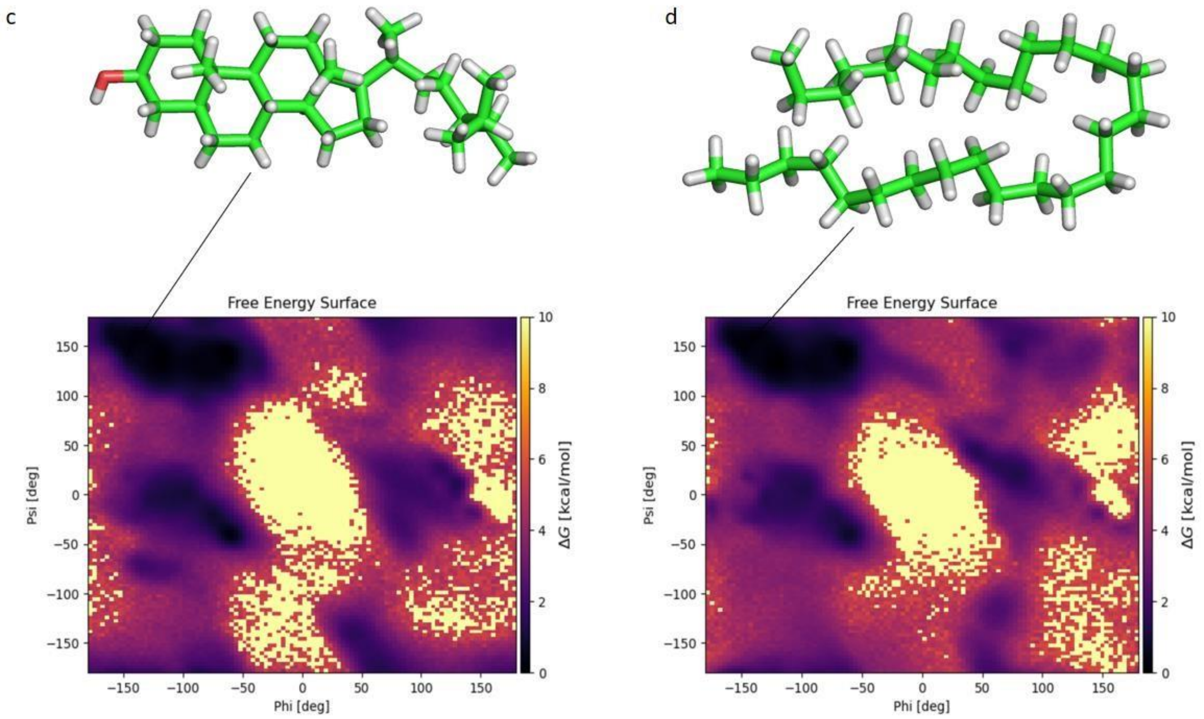


Fig S2: Free energy Surface Plot for the respective Neurotrophins along with the structure of the ligand at the most stable energy state where x and y axis represents the phi and psi angles and the color bar corresponds to the free energy change in kcal/mol. (a) Free energy surface plot of BDNF showing the most stable region in terms of phi-psi angle and the corresponding structure of the ligand (Vitamin E) (b) Free energy surface plot of NT3 along with the corresponding ligand structure (BPA) (c) Free energy surface plot of NT4 along with the corresponding structure of ligand (Stigmasterol) (d) Free energy surface plot of NGF along with the corresponding structure of ligand (Nonacosane)

**REFERENCES**

Abraham, M. J., Murtola, T., Schulz, R., Páll, S., Smith, J. C., Hess, B., & Lindah, E. (2015). Gromacs: High performance molecular simulations through multi-level parallelism from laptops to supercomputers. *SoftwareX*, *1–2*, 19–25. https://doi.org/10.1016/j.softx.2015.06.001

Ajay., Bemis, G. W., & Murcko, M. A. (1999). Designing Libraries with CNS Activity. *J. Med. Chem*. 42, 24, 4942-4951. https://doi.org/10.1021/jm990017w

Allinger N. L. (1977). Conformational Analysis. 130. MM2. A Hydrocarbon Force Field Utilizing V1 and V2 Torsional Terms. *Journal of the American Chemical Society*, 99 (25), 81278134. doi:10.1021/ja00467a001.

Ames, B. N., Gurney, E. G., Miller, J. A., & Bartsch, H. (1972). Carcinogens as Frameshift Mutagens: Metabolites and Derivatives of 2-Acetylaminofluorene and other Aromatic Amines Carcinogen. *Proc Natl Acad Sci USA*, 69(11):3128-3132. https://doi.org/10.1073/pnas.69.11.3128

Duhovny, D., Nussinov, R., & Wolfson, H. J. (2002). Efficient unbound docking of rigid molecules. Lecture Notes in Computer Science (Including Subseries Lecture Notes in Artificial

Intelligence and Lecture Notes in Bioinformatics), 2452, 185–200. https://doi.org/10.1007/3540-45784-4_14

Schneidman-Duhovny, D., Inbar, Y., Nussinov, R., & Wolfson, H. J. (2005). PatchDock and SymmDock: servers for rigid and symmetric docking. Nucl. Acids. Res. 33: W363-367.

Yamashita, S., Furubayashi, T., Kataoka, M., Sakane, T., Sezaki H., & Tokuda, H. (2000). Optimised conditions for prediction of intestinal drug permeability using Caco-2 cells. *Eur J Pharm Sci*, 10(3):195-204. https://doi.org/10.1016/s0928-0987(00)00076-2

Zhao, Y. H., Le, J., Abraham, M. H., Hersey, A., Eddershaw, P. J., Luscombe, C. N., Butina, D., Beck, G., Sherbone, B., Cooper, I., & Platts, J. A. (2001). Evaluation of human intestinal absorption data and subsequent derivation of a quantitative structure-activity relationship (QSAR) with the Abraham descriptors. *J Pharm Sci.*, 90(6):749-84. https:doi.org/10.1002/jps.1031
